# Robust functional ultrasound imaging in the awake and behaving brain: a systematic framework for motion artifact removal

**DOI:** 10.1101/2025.06.16.659882

**Authors:** Samuel Le Meur-Diebolt, Felipe Cybis Pereira, Jean-Charles Mariani, Andrea Kliewer, Miguel Farinha-Ferreira, Adrien Bertolo, Bruno-Félix Osmanski, Zsolt Lenkei, Thomas Deffieux

## Abstract

Functional ultrasound imaging (fUSI) is a promising tool for studying brain activity in awake and behaving animals, offering insights into neural dynamics that are more naturalistic than those obtained under anesthesia. However, motion artifacts pose a significant challenge, introducing biases that can compromise the integrity of the data. This study provides a comprehensive evaluation and benchmarking of strategies for detecting and removing motion artifacts in transcranial fUSI acquisitions of awake mice. We evaluated 792 denoising strategies across four datasets, focusing on clutter filtering, scrubbing, frequency filtering, and confound regression methods. Our findings highlight the superior performance of adaptive clutter filtering and aCompCor confound regression in mitigating motion artifacts while preserving functional connectivity patterns. We also demonstrate that high-pass filtering is generally more effective than band-pass filtering in the presence of motion artifacts. Additionally, we show that with effective clutter filtering, scrubbing may become optional, which is particularly beneficial for experimental designs where motion correlates with conditions of interest. Based on these insights, we propose four optimized denoising paradigms tailored to different experimental constraints, providing practical recommendations for enhancing the reliability and reproducibility of fUSI data. Our findings challenge current practices in the field and have immediate practical implications for existing fUSI analysis workflows, paving the way for more sophisticated applications of fUSI in studying complex brain functions and dysfunctions in awake experimental paradigms.

## 1 Introduction

Functional ultrasound imaging (fUSI) has the potential to become a useful and cost-effective tool for studying brain activity in awake and behaving mice (Brunner et al., 2021; Tiran et al., 2017), the main animal model for preclinical research (Soufizadeh et al., 2024). Awake imaging is one of the most promising applications of fUSI, enabling studies of brain function in a more naturalistic context than anesthetized paradigms. While anesthetized preparations allow for simpler experimental setups, lower animal stress, and reduced motion artifacts, they often fail to reproduce the complex dynamics of the conscious brain. Indeed, anesthetics have been shown to fundamentally alter neural activity, neurovascular coupling (NVC), and functional networks (Barttfeld et al., 2015; Gao et al., 2017; Pisauro et al., 2013). Additionally, awake setups allow studying cognitive processes and behavior, which are essential to understand normal brain function and model neurological disorders (Ferenczi et al., 2016; Tsurugizawa et al., 2020). Proof of concept studies in awake rodents have already demonstrated the use of fUSI to study awake functional connectivity (FC) (Ferrier et al., 2020), locomotion (Bergel et al., 2020), pharmacological alterations (Rabut et al., 2020; Vidal et al., 2020a; Vidal et al., 2020b), and stroke (Brunner et al., 2023).

fUSI shares several similarities with functional magnetic resonance imaging (fMRI) when it comes to imaging the brain of awake mice. Both techniques rely on the NVC to measure local neural activity (Mace et al., 2013; Ogawa et al., 1990): fMRI measures the blood-oxygen-level-dependent (BOLD) signal, while fUSI measures the power Doppler signal where voxel intensity is proportional to the cerebral blood volume (CBV). Both generally use surgically implanted head-posts to immobilize the mouse head relative to the sensor. Importantly, both are biased by animal motion, even after head-fixation and animal habituation (Ferrier et al., 2020; Mandino et al., 2023). Indeed, animal motion leads to large variations in both BOLD and power Doppler signals, referred to as “motion artifacts”—also known as “flash” or “clutter” artifacts in ultrasound imaging. These motion artifacts are particularly problematic in studies of signal covariance between brain regions—referred to as “functional connectivity (FC)”—, as they lead to spurious correlations (Power et al., 2012). However, while motion artifacts have been extensively studied in fMRI (Friston et al., 1996; Hajnal et al., 1994; Parkes et al., 2018; Power et al., 2012; 2014; Satterthwaite et al., 2013; 2017; Van Dijk et al., 2012), they have yet to receive the same attention in the fUSI literature.

Notably, the origin of motion artifacts has not yet been discussed in the context of fUSI. In particular, animal motion is known to trigger widespread neuronal activity (Steinmetz et al., 2019) and could therefore lead to large and relevant increases of CBV through the NVC. However, the extreme magnitude of power Doppler surges seen during motion artifacts— often exceeding 400% above baseline in some areas—cannot be explained by neural activity alone. While the lack of interest for these artifacts can be partly explained by the relative novelty of fUSI compared to fMRI—the first fUSI studies having been published two decades after the seminal fMRI article (Belliveau et al., 1991; Macé et al., 2011)—, the confounding effect of motion artifacts coupled with the rapid gain in popularity of fUSI in the neuroscience community calls for immediate attention.

Although head-fixation setups generally prevent visible brain movements in transcranial fUSI acquisitions, even sub-pixel brain movements can trigger significant power Doppler surges across the entire field of view (FOV). Unfortunately, the high spatial frequencies of power Doppler volumes and the nature of these artifacts make traditional framewise realignment strategies ineffective. To date, no fUSI study has successfully implemented framewise realignment to correct motion artifacts in awake head-fixed rodents. Instead, most fUSI studies use basic denoising techniques borrowed from fMRI to mitigate motion artifacts and other non-neuronal noise sources. These include simple thresholding to remove high-motion time points—a method referred to as scrubbing (Power et al., 2012)—, global signal regression (GSR), and band-pass filtering. These approaches are applied with minimal adaptation to the specific characteristics of fUSI data (Ferrier et al., 2020). Unfortunately, these methods come with trade-offs. First, scrubbing periods of animal movement cannot be used in behavioral paradigms where motion is expected to correlate with conditions of interest. Furthermore, standardized movement-detection methods have yet to be introduced and validated for fUSI. Second, the use of GSR is controversial in the fMRI literature, as it can remove genuine neural signals that are spatially distributed across the brain and lead to negative correlations that are difficult to interpret (Murphy & Fox, 2017). These concerns are even more pronounced in fUSI, where the FOV is generally restricted to narrow brain slices, making the “global signal” not truly global. While more advanced confound regression methods have been proposed in the fMRI literature—for example, CompCor (Behzadi et al., 2007), FIX (Griffanti et al., 2014), ICA-AROMA (Pruim et al., 2015)—, they have not yet been evaluated on fUSI data. Finally, low-pass filtering is often used to retain low-frequency signals of interest thought to be related to neural activity, while removing high-frequency noise—for example, cardiac, respiratory (Fox & Raichle, 2007). However, low-pass filters inflate temporal autocorrelations, leading to spurious correlations in the presence of motion artifacts (Ebisuzaki, 1997).

Rather than addressing motion artifacts in the processed power Doppler data, technical development has mostly focused on improving clutter filters to remove unwanted clutter signals from the beamformed ultrasound signals (Baranger et al., 2018; Demene et al., 2015; Song et al., 2017). Indeed, the main challenge in power Doppler imaging stems from the mixture of signals in the recorded radio frequency (RF) data: stationary or slow-moving clutter (for example, skull, tissue, vessel walls) must be separated from red blood cell scatterers that constitute the signal of interest. Singular value decomposition (SVD) emerged as a powerful approach to this separation problem, surpassing traditional frequency and regression filters by exploiting the distinct properties of clutter and scatterer signals (Ledoux et al., 1997; Yu et al., 2007). The use of SVD for clutter filtering is motivated by the observation that clutter areas reflect highly energetic echoes and are generally correlated both spatially and temporally. In contrast, blood scatterers reflect less energy and exhibit lower spatial and temporal correlation. Thus, singular vectors with the highest singular values are likely to correspond to clutter components, and may be filtered out.

Soon after the seminal article on fUSI by Macé and colleagues (Macé et al., 2011), SVD-based clutter filtering was shown to be particularly effective in suppressing clutter signals for ultrafast power Doppler imaging (Demene et al., 2015). Since then, efforts have been made to compare estimators of the threshold between clutter and scatterer components in SVD clutter filters (Baranger et al., 2018; Song et al., 2017). Despite these methodological advances, the vast majority of fUSI studies in awake rodents continue to use static thresholds for SVD-based filters, typically removing a fixed number of singular vectors based on empirical observations of the resulting power Doppler images (Brunner et al., 2021; Ferrier et al., 2020; Nunez-Elizalde et al., 2022; Rabut et al., 2020). However, this static approach is known to be suboptimal for awake imaging, as the boundary between clutter and scatterer components fluctuates substantially during acquisition due to animal motion (Baranger et al., 2018). On the other hand, while adaptive approaches may more effectively remove clutter during motion periods, they can paradoxically remove too much signal energy during high-motion periods, producing an apparent drop in global CBV. This contradicts the expected physiological response, where increased movement would typically be associated with increased brain perfusion.

Consequently, there is a complex interplay between signal processing choices and physiological interpretation in fUSI—a challenge that requires careful consideration when designing analyses and interpreting results. In this work, we first characterize the motion artifacts in transcranial fUSI acquisitions of the awake mouse brain and provide a set of spatial and temporal quality control (QC) metrics to identify acquisitions with high motion artifacts. Next, we benchmark the performance of several denoising strategies, quantifying their sensitivity to motion artifacts as well as their capacity to preserve relevant FC patterns. To ensure the robustness of our findings, we evaluate these strategies across three different datasets. Finally, we develop practical recommendations for optimizing fUSI data denoising in different experimental contexts.

## 2 Methods

### 2.1 Datasets

This study re-used 4 datasets of transcranial fUSI acquisitions from awake head-fixed mice, comprising a total of 193 male C57BL/6J mice (see Table 1). Dataset 1 represents a subset of data previously published in (Mariani et al., 2024). Datasets 2 and 3 contain subsets of data acquired for articles in preparation, with similar protocol as dataset 1. Dataset 4 represents a subset of data previously published in (Bertolo et al., 2023). All acquisitions consist of 20 to 30 minutes of awake and behaving condition. Details on surgical procedures and imaging preparation can be found in (Mariani et al., 2024) and (Bertolo et al., 2023). All experimental procedures were conducted with approval from the relevant institutional ethical authorities.

**Table 1:** Datasets characteristics. The number of mice, acquisitions per mice (minimum and maximum), total acquisitions, and average brain axial velocity are shown for each dataset.

| | FOV | $N_{\text{mice}}$ | $N_{\text{acquisitions}}$ | $N_{\text{mice post-QC}}$ | $N_{\text{acquisitions post-QC}}$ | Axial velocity mean ± SD (mm/s) | Sampling rate (Hz) | Duration (min) |
| --- | --- | --- | --- | --- | --- | --- | --- | --- |
| Dataset 1 | 2D | 28 | 120 | 25 | 105 | $(3.64 \pm 1.06) \cdot 10^{-2}$ | 2.5 | 20 |
| Dataset 2 | 2D | 89 | 122 | 84 | 117 | $(2.91 \pm 0.49) \cdot 10^{-2}$ | 2.5 | 30 |
| Dataset 3 | 2D | 70 | 314 | 65 | 288 | $(2.62 \pm 0.79) \cdot 10^{-2}$ | 2.0 | 20 |
| Dataset 4 | 3D | 6 | 6 | 6 | 6 | $(1.98 \pm 1.51) \cdot 10^{-2}$ | 0.42 | 20 |

### 2.2 Functional ultrasound imaging

Datasets 1 to 3 were acquired using a prototype fUSI system (Inserm Biomedical Ultrasound ART & Iconeus, Paris, France) with a 15 MHz ultrasound probe (128 elements, 110 µm pitch, 8 mm elevation focal distance, 400 µm elevation focal width). Dataset 4 was acquired using an Iconeus One fUSI system (Iconeus, Paris, France) with a 15.625 MHz multiarray probe (IcoPrime-4D Multi-array, 110 µm pitch, 8 mm elevation focal distance, 500 µm elevation focal width, Iconeus, Paris, France). For all datasets, probes were mounted on a motorized stage, allowing for translations along three axes (1 µm precision) and rotations along the vertical axis (0.1° precision). For dataset 1 to 3, the 2D plane of interest was selected visually based on anatomical vascular landmarks (Mariani et al., 2024). The 2D ultrafast acquisition sequence (datasets 1 to 3) consisted of 11 titled plane waves ((−10 to 10)° with 2° step) transmitted at a pulse repetition frequency (PRF) of 5.5 kHz, leading to a 500 Hz sampling rate after beamforming and compounding. The multiarray ultrafast sequence consisted of 4 × 8 tilted plane waves ((−12 to 12)° with 3.43° step) transmitted at a PRF of 4 kHz, leading to a 500 Hz sampling rate after beamforming and compounding. Acquisitions from dataset 3 included a 100 ms delay after each set of 200 compound volumes.

### 2.3 Axial velocity estimation

Axial velocity was estimated from the beamformed in-phase/quadrature (I/Q) signals using the Kasai autocorrelation technique (Kasai et al., 1985), from which we will give a brief overview.

Let IQ(*x, y, z, t*) be the complex-valued beamformed I/Q signal at voxel (*x, y, z*) and time point *t*. IQ can be written as

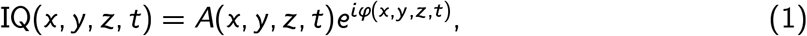

where *A*(*x, y, z, t*) is the time-varying amplitude and φ(*x, y, z, t*) is the time-varying phase. The mean velocity of the scatterers along the vertical axis *v*_*z*_ can then be estimated according to

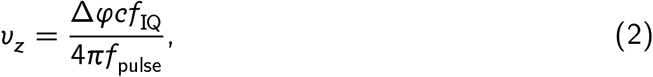

where Δφ is the phase shift between two adjacent samples, *f*_0_ is the frequency of the transmitted ultrasound pulse, *c* is the speed of sound, and *f*_IQ_ is the sampling frequency of the beamformed I/Q signals.

Now, remark that we can compute the phase shift as

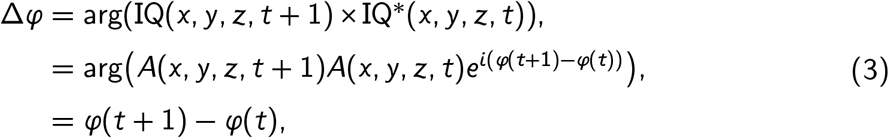

where arg is the complex argument and IQ^∗^ denotes the complex conjugate of IQ. Thus, we can estimate *v*_*z*_ as

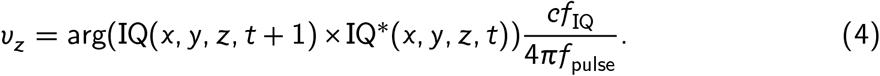

In practice, this velocity estimator is often noisy. Estimation was improved by filtering beamformed I/Q signals before velocity estimation using a Butterworth low-pass filter with a cutoff frequency of 30 Hz and order 6. The resulting velocity volumes were then smoothed using a spatial median filter with a kernel of 3 × 3 × 3 voxels.

Finally, absolute axial velocity acquisitions were obtained with same sampling rate as the power Doppler acquisitions by computing and averaging the absolute value of the velocity across blocks of 200 compound I/Q volumes, or 400 ms.

### 2.4 Clutter filtering and power Doppler integration

Before power Doppler integration, each block of 200 compound volumes (that is, 400 ms) was processed using a singular value decomposition (SVD)-based clutter filter to remove clutter signals (Demene et al., 2015). SVD-based clutter filters rely on the assumption that clutter signals have higher energy and spatiotemporal correlation than blood signals. Thus, clutter signals can be identified as temporal singular vectors with high singular values and regressed out.

Let *X* be a (time, voxels) matrix corresponding to a block of compound volumes. To avoid biases from outlier signals, a mask may be used to select only a subset of voxels in *X*. Let *M* be a (time, mask) matrix corresponding to such a subset, from which we will compute clutter signals. In practice, clutter signals are computed using the eigenvalue decomposition (EVD) of the left Gram matrix *MM*^*H*^ (where *M*^*H*^ is the conjugate transpose of *M*):

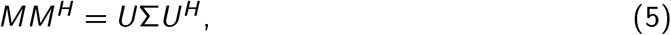

where columns of *U* are the eigenvectors of *MM*^*H*^ (or equivalently the temporal singular vectors of *M*), and Σ is the diagonal matrix of eigenvalues (or equivalently the squared singular values of *M*). Note that since *U* is unitary—that is, *U* ^*H*^*U* = *U U* ^*H*^ = *I* where *I* is the identity matrix—, it follows that eigenvalues correspond to the energy of their corresponding eigenvectors. Thus, clutter signals are selected as a subset of eigenvectors *U*_clutter_ with highest eigenvalues—that is, highest energy—and regressed out from *X*:

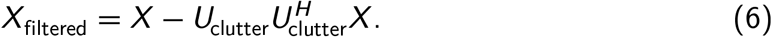

If we denote *M*_filtered_ the (time, mask) matrix corresponding to the same subset of voxels as *M* in *X*_filtered_, it follows that the energy of *M*_filtered_ (defined as its Frobenius norm) is equal to the sum of eigenvalues that were not retained in the clutter subspace.

In this study, we compared three methods to select clutter signals:

▪ The “static SVD*T* “clutter filter selects a fixed number *T* of clutter signals with highest eigenvalues from each compound block. The threshold *T* is constant across all compound blocks.
▪ The “static SVD*T* + high-pass” clutter filter applies the static-SVD clutter filter with threshold *T*, followed by a Butterworth high-pass filter with a cutoff frequency of 20 Hz and order 8. Given the ultrasound frequency of 15.625 MHz and assuming a speed of sound of approximately 1 540 m s^−1^, this high-pass cutoff corresponds to retaining blood velocities above ≈ 1 mm s^−1^ (Mace et al., 2013), just above the typical range of velocities in capillaries and cortical ascending venules (Chaigneau et al., 2003; Nguyen et al., 2011).
▪ The “constant-energy SVD*T* “clutter filter selects a variable number of clutter signals such that the energy—or in other words, the sum of retained singular values—of *M*_filtered_ approaches but does not exceed a threshold *C*. For better comparison with static SVD filters, the energy threshold *C* is computed for each acquisition as the minimum energy of *M*_filtered_ observed across all compound blocks, when filtered with a static-SVD clutter filter with threshold *T*.

We benchmarked the performance of these methods using three thresholds *T* : 20, 40, and 60. Additionally, we used either the whole FOV or a brain mask to compute clutter signals.

After clutter filtering, power Doppler volumes were computed from each block of 200 compound volumes by averaging their squared modulus across time. The resulting sampling rates were 2.5 Hz for datasets 1 and 2, and 2.0 Hz for dataset 3.

### 2.5 Atlas registration and resampling

For each of datasets 1, 2, and 3, we generated a separate custom power Doppler template through iterative affine registration. The following procedure was repeated for each dataset. First, we selected only the power Doppler acquisitions computed using the static SVD60 clutter filter without brain mask. From these acquisitions, we computed average and decibel-scaled volumes. The initial template iteration starts as the average dataset volume. Then, each decibel-scaled volume is registered and resampled to the current template iteration. The template is updated after each iteration by averaging all resampled volumes. We found that two iterations were sufficient to provide a satisfactory template. This process is similar to that used in the RABIES software and for creation of the Allen Mouse Brain Common Coordinate Framework (CCFv3) template (Desrosiers-Grégoire et al., 2024; Wang et al., 2020).

Registration was performed using Python 3.12 and the SimpleITK Python package version 2.4.0 (Yaniv et al., 2017). Affine transformations were initialized at center of mass. Optimization was performed using the normalized cross-correlation metric and the gradient descent optimizer with a maximum of 1000 iterations, a minimum convergence value of 10^−6^, a convergence window of size 10, and automatic estimation of the learning rate after each iteration.

Once each custom template was generated, it was manually masked to remove non-brain voxels and registered to a standard power Doppler template using an affine transform with mutual information metric and otherwise same parameters as above. The standard template was obtained from (Nouhoum et al., 2021) and was already aligned to the Allen Mouse Brain CCFv3.

Dataset 4 was already registered to the Allen Mouse Brain CCFv3 template (Bertolo et al., 2023). Masks of brain, white matter, and cerebrospinal fluid (CSF) voxels were obtained for each dataset from the Allen Mouse Brain CCFv3 atlas (Wang et al., 2020).

Finally, all axial velocity and power Doppler acquisitions were resampled to their respective custom template for datasets 1 to 3, or to the first acquisition for dataset 4. For datasets 1 to 3, resampling was performed using the affine transformation obtained from the static SVD60-filtered acquisition at the last iteration of the template generation. For dataset 4, resampling was performed using the existing registration transforms. Note that 3D+t acquisition from dataset 4 were slice-timing corrected using linear interpolation prior to resampling. Linear interpolation was preferred due to intensity artifacts observed when using higher interpolation orders. This led to datasets of aligned axial velocity and power Doppler acquisitions with corresponding Allen Mouse Brain CCFv3 coordinates.

### 2.6 Spatial and temporal characteristics of motion artifacts

To better understand the nature of motion artifacts in fUSI, we investigated their spatial and temporal characteristics Figure 1.

**Figure 1.**
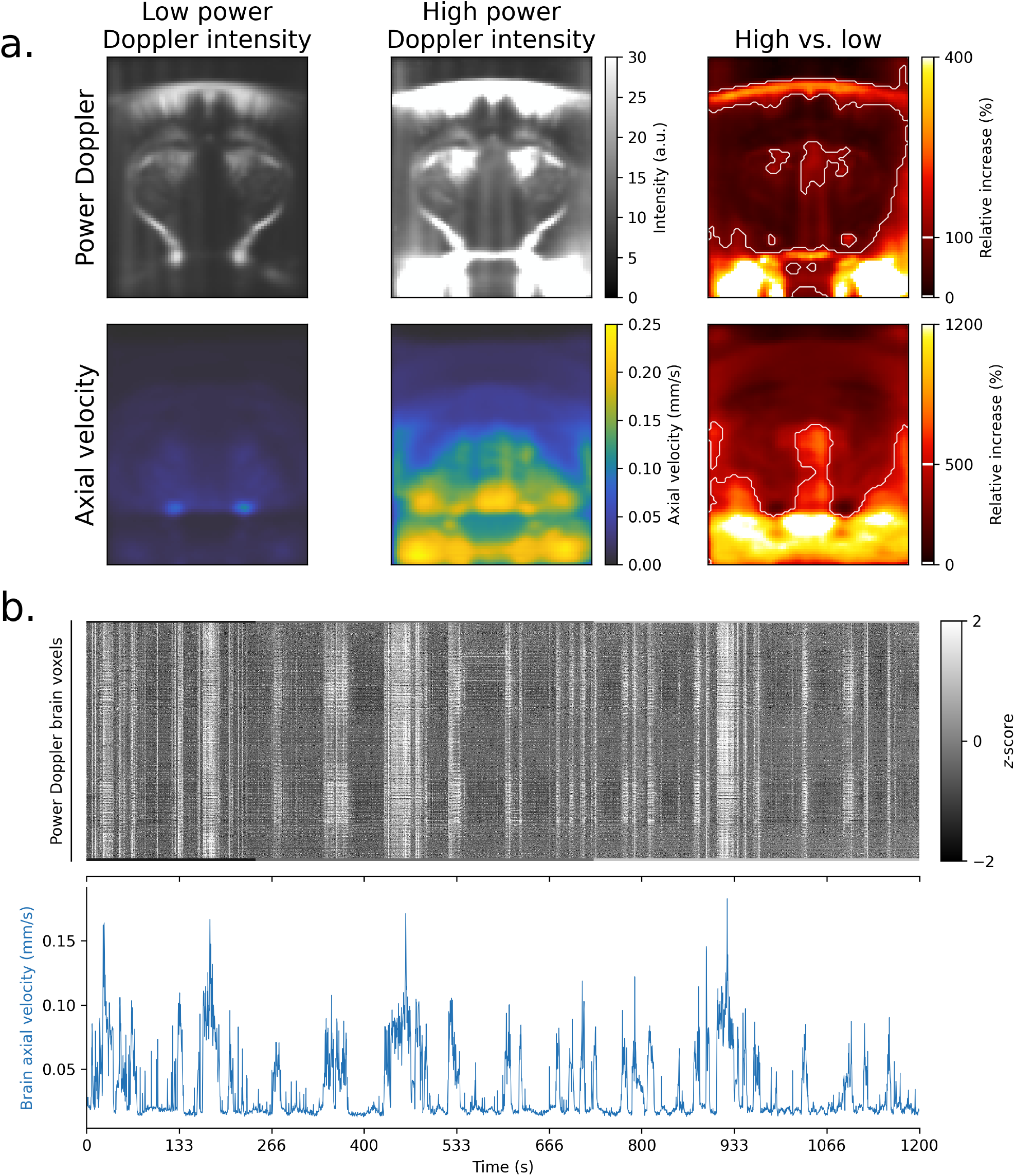
Motion artifacts show distinct spatial patterns with large and synchronous power Doppler surges across the whole FOV. (**a**.) Average power Doppler and axial velocity frames across dataset 1 during high vs. low power Doppler intensity periods. Low- and high-intensity frames were selected as the 5% lowest, respectively highest, average power Doppler intensity frames. High-intensity power Doppler frames show large surges in inferior cerebral veins, around the skull, and in lateral areas closer to muscles. In contrast, axial velocity frames show increased velocities in ventral brain regions and areas below the brain, while the cortex and dorsal skull areas show lower velocities. The lower part of the skull is particularly visible in the relative increase. (**b**.) Carpet plot of standardized power Doppler time series from an example acquisition of dataset 1 (**top**) and corresponding global axial velocity (GV) (**bottom**). Large artifacts appear simultaneously across all brain voxels during high velocity periods.

First, we generated average power Doppler and axial velocity images to compare low- and high-intensity periods. For each acquisition, we selected the top 5% of power Doppler frames with lowest, respectively highest, intensity, and computed the average power Doppler volume across these frames. We then computed the relative difference between these average volumes. The average velocity volumes and their relative difference was computed for corresponding axial velocity frames, so as to compare axial velocities between low- and high-intensity power Doppler frames. Finally, we computed group-level averages of power Doppler, axial velocity, and their relative differences across all acquisitions.

Then, we generated a “carpet plot” (Power et al., 2012) of an example acquisition from Dataset 1 by displaying standardized (mean centered and scaled to unit variance) power Doppler time series from brain voxels in gray scale. This visualization allows to identify the presence of spatially correlated signals.

Finally, we investigated the temporal characteristics of motion artifacts by modeling power Doppler signals against the average axial velocity signal from brain voxels. To this end, we used a finite impulse response (FIR) model where the power Doppler signal at each voxel was modeled as a linear combination of all delayed versions of the average axial velocity signal between −12 s and 12 s. Subject-level FIR models were fitted for each mouse using the first-level general linear model (GLM) class of the Nilearn Python package (nilearn.glm.first_level.FirstLeveLModel). A second-level GLM was then fitted to the resulting first-level beta-maps using the second-level GLM class of Nilearn (nilearn.glm.second_level.SecondLevelModel). Resulting beta-maps were converted to *z*-scores, and average *z*-scores were computed across three regions of interest (ROIs): the primary somatosensory cortex, hippocampus, and thalamus.

### 2.7 Quality control metrics

We computed a set of 3 spatial and 3 temporal quality control (QC) metrics for each power Doppler acquisition.

▪ Spatial QC metrics consisted of the average power Doppler intensity, the temporal coefficient of variation (CV), and the average axial velocity. The average power Doppler intensity and average axial velocity were computed by temporally averaging the respective volumes, while the temporal CV was computed as the ratio of the temporal standard deviation of the power Doppler intensity to its temporal average.
▪ Temporal QC metrics consisted of the relative global power Doppler signal (rGS), global axial velocity (GV), and relative DVARS (rDVARS).
  ▸ The global power Doppler signal (GS) was computed by averaging time series from a power Doppler acquisition across brain voxels. The rGS was then computed as

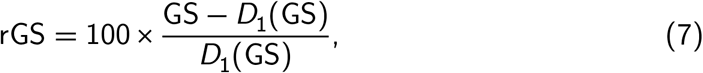

where *D*_1_(GS) is the first decile of GS, used as a robust measure of baseline.
  ▸ The GV was computed by averaging time series from an axial velocity acquisition across brain voxels.
  ▸ The rDVARS signal was computed following recommendations by (Nichols, 2017). DVARS measures the rate of change of signals across the imaged brain volume, at each time point (Power et al., 2012; Smyser et al., 2010). DVARS is computed by differentiating the voxel time series (using backward differences), then computing the root mean square (RMS) signal change over brain voxels:

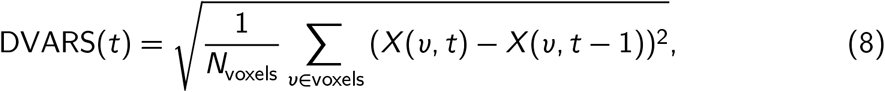

where *X*(*v, t*) is the time series at voxel *v* and time point *t* and *N*_voxels_ is the number of voxels. Then, the rDVARS signal was computed by scaling DVARS by its null mean µ_0_,

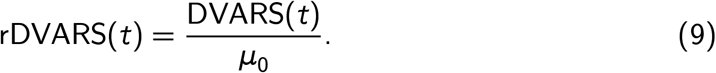

Assuming an autoregressive order 1 (AR(1)) model of temporal noise with parameter ρ_*v*_, µ_0_ can be estimated via

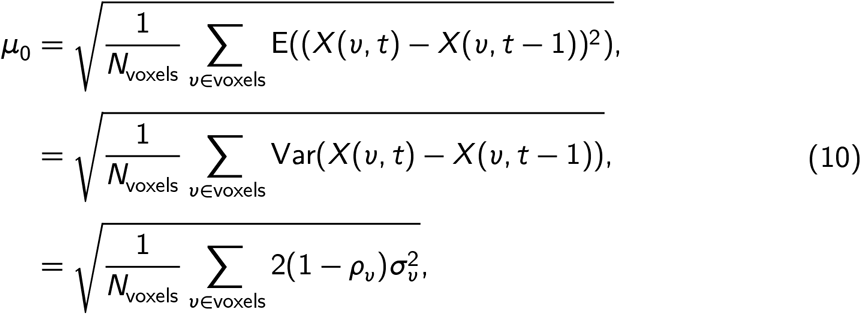

where σ_*v*_ is the temporal standard deviation at voxel *v* and can be robustly estimated usign the interquartile range (IQR),

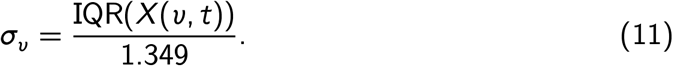

### 2.8 Core denoising pipeline

The core FC denoising pipeline consisted of spatial smoothing, scrubbing, frequency filtering, and confound regression (see Figure 4 a.). The FC denoising pipeline follows standards from the fMRI literature (Lindquist et al., 2019; Power et al., 2014). Steps are applied in the following sequence:

1. **Spatial smoothing:** spatial smoothing was systematically applied with a Gaussian kernel of 0.3 mm full width at half maximum (FWHM).
2. **Scrubbing:** scrubbing was optionally applied before any other temporal denoising step to avoid signal surges that may bias frequency filtering, and confound regression. Since the frequency filter requires regular sampling, scrubbing was performed by replacing high motion volumes by the average signal from non-motion volumes. We benchmarked the use of 3 metrics to identify high-motion volumes: relative global power Doppler signal (rGS), relative DVARS (rDVARS), and global axial velocity (GV). For each metric, we used a single strict threshold: 10% for rGS, 2.0 for rDVARS, and 0.01 mm s^−1^ above the baseline (estimated using the first decile) for GV. These thresholds were selected based on visual inspection of the signals to obtain a strict-thresholding procedure. Note that any acquisition with less than 5 minutes retained after scrubbing was discarded.
3. **Frequency filtering:** we benchmarked the use of either a high-pass filter with a cutoff frequency of 0.01 Hz or a band-pass filter with cutoffs (0.01 to 0.1) Hz. Both high-pass and band-pass filters were implemented using a forward-backward Butterworth filter of order 6. This filter design was selected to avoid phase distortion. After frequency filtering, the previously replaced high-motion volumes in step 2 were removed entirely.
4. **Confound regression:** we benchmarked the use of 5 methods to compute confounds regressors:

▪ *Global brain signal*, computed as the average power Doppler signal over brain voxels,
▪ *tCompCor* (Behzadi et al., 2007), computed as the first 5 principal components of the power Doppler signals over the 5% voxels with highest temporal standard deviation,
▪ *aCompCor* (Behzadi et al., 2007), computed as the first 5 principal components of the power Doppler signals over white matter and CSF voxels (representing approximately 16% of the brain volume in datasets 1 to 3, and approximately 10% in dataset 4),
▪ *Random CompCor*, computed as the first 5 principal components of the power Doppler signals over a random voxel mask. The random mask was computed by randomly selecting as many voxels in the brain as there are white matter and CSF voxels. One random mask was generated for each dataset.
▪ *Low-variance CompCor*, computed as the first 5 principal components of the power Doppler signals over voxels with lowest temporal standard deviation. As many voxels as the aCompCor mask are selected.

Additionally, we also benchmarked a *no confound regression* method, where no confounds were regressed out. Following recommendations by Lindquist and colleagues, confounds underwent the same scrubbing and frequency filtering steps as power Doppler signals (Lindquist et al., 2019). Confound regression was performed by modeling voxel signals using ordinary least squares (OLS) linear regression and removing the contribution of confounds from the signals,

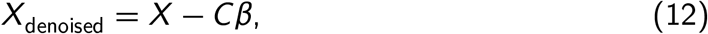

where *X* is the (time, voxels) matrix of power Doppler signals, *C* is the (time, confounds) confounds matrix, and β is the (confounds, voxels) matrix of OLS coefficients.

Combining all clutter filter and FC denoising parameters leads to 864 denoising strategies. Note however that the constant-energy SVD*T* clutter filter strongly impacts the global power Doppler signal (GS) such that rGS-based scrubbing becomes ineffective. We therefore excluded these combinations from our analysis *a priori*, resulting in 792 unique strategies.

The core FC denoising pipeline was adapted from the nilearn.signal.clean function from Nilearn 0.11.0 (Nilearn contributors, 2024), with adaptations to the scrubbing step to remove the cubic spline interpolation, which introduces large biases when interpolating long periods of high-motion volumes.

### 2.9 Standard denoising strategy

The standard denoising strategy, used as ground-truth for the FC similarity metric, was selected from common practices identified across recent fUSI studies (Bertolo et al., 2023; Ferrier et al., 2020; Rabut et al., 2020). It consisted of a static SVD60 clutter filter without brain mask, scrubbing using rGS above 10%, [0.01, 0.1] Hz band-pass filtering, and GSR. We visually confirmed that this strategy provided good quality group-level seed-based maps in all datasets.

### 2.10 Seed-based functional connectivity

We computed seed-based FC maps following FC denoising. Seeds were placed in the primary somatosensory cortex, hippocampus, and thalamus, in both the left or right hemisphere, leading to 6 seed-based maps per denoised acquisition. Seed ROIs were computed as spheres of radius 0.3 mm centered on the corresponding Allen Mouse Brain CCFv3 ROIs. Seed signals were then computed from denoised acquisitions as the average power Doppler signal over all voxels in the seed sphere. Seed-based maps were finally computed using the Pearson correlation coefficient between seed signals and all other brain voxels. Average seed-based maps were computed by first transforming correlation coefficients to Fisher’s *z*, averaging the *z* values, then backtransforming to correlation coefficients (Alexander, 1990).

### 2.11 Motion robustness metric

To evaluate the robustness of the denoising strategies (excluding scrubbing) to motion artifacts, we introduced a motion robustness metric. The motion robustness metric indexes how much a given denoising strategy is susceptible to motion artifacts. For each set of clutter filter, frequency filter, and confound regression parameters, we computed the mean squared error (MSE) between seed-based maps obtained without scrubbing and seed-based maps obtained with global axial velocity (GV)-based scrubbing. Then, the motion robustness metric was computed for each dataset and each set of strategies as the average MSE across mice. To avoid biases from the variable number of acquisition per mouse, averaging was performed across acquisitions of each mouse, then across mice. The standard deviation of the MSE was computed across mice to evaluate the variability of the metric. Since scrubbing may lead to discarded acquisitions if less than 5 minutes of data are retained, we only considered denoising strategies that retain at least two thirds of the mice in each dataset. Thus, each set of clutter filter, frequency filter, and confound regression parameters was indexed by an average MSE and a standard deviation of the MSE. The combination of all denoising parameters excluding also scrubbing lead to 216 denoising strategies.

### 2.12 Functional connectivity similarity metric

To evaluate the relevance of the denoising strategies for functional connectivity (FC) analyses, we introduced an FC similarity metric. The FC similarity metric indexes how much a denoising strategy is able to preserve group-level FC patterns in each acquisition. The “ground-truth” group-level FC patterns were defined as the average seed-based maps across mice obtained using the standard denoising strategy. For each set of denoising parameters, we first computed binary masks of correlated regions by thresholding seed-based maps to keep 10% of voxels with highest absolute correlation values. Then, we computed the Dice coefficient between each binary mask and the binary mask obtained from the group-level FC patterns using the same thresholding. The FC similarity metric was computed as the average Dice coefficient across mice. To avoid biases from the variable number of acquisition per mouse, averaging was performed across acquisitions of each mouse, then across mice. The standard deviation of the Dice coefficient was computed across mice to evaluate the variability of the metric. Since scrubbing may lead to discarded acquisitions if less than 5 minutes of data are retained, we only considered denoising strategies that retain at least two thirds of the mice in each dataset. Thus, each set of denoising parameters was indexed by an average Dice coefficient and a standard deviation of the Dice coefficient.

### 2.13 Computation and visualization

All computations and visualizations were performed using a proprietary Python package (Iconeus, France) developed using Python 3.12 and the following dependencies: numpy 2.1.2 (Harris et al., 2020), scipy 1.14.1 (Virtanen et al., 2020), scikit-learn 1.5.2 (Pedregosa et al., 2011), nibabel 5.3.1 (Brett et al., 2024), nilearn 0.11.0 (Nilearn contributors, 2024), nitime 0.11.0 (Rokem et al., 2009), pybids 0.18.1 (Yarkoni et al., 2019), matplotlib 3.9.2 (Hunter, 2007), seaborn 0.13.2 (Waskom, 2021), statsmodels 0.14.2 (Seabold & Perktold, 2010), and pandas 2.2.3 (The pandas development team, 2020).

## 3 Results

### 3.1 Motion artifacts exhibit distinct spatial patterns and delayed effects despite head fixation

In fUSI, movements of the imaged tissues relative to the probe cause significant power Doppler signal increases across large portions of the field of view (FOV). These surges, often called “flash artifacts” or “motion artifacts”, occur despite head fixation in awake mice and significantly bias correlation-based analyses—broadly defined as functional connectivity (FC).

Motion artifacts show distinct spatial patterns both on power Doppler and axial velocity images (Figure 1 a.). Comparing frames with high vs. low average power Doppler intensity, we observed that increases are primarily located in inferior cerebral veins, around areas of strong reflectivity (for example, the skull), and in lateral areas closer to muscles. Additionally, the cortex and thalamus show stronger power Doppler increases, while the hippocampus and some deeper brain structures are less impacted. In contrast, corresponding axial velocity frames show non-uniform distributions of velocities across the brain, with higher velocities in ventral brain regions and areas below the brain, and lower velocities in the cortex and dorsal skull areas. This pattern likely results from the mechanical constraints of the head-fixed setup, where the top of the skull is secured to a metal plate.

Unsurprisingly, frames with high power Doppler intensity are associated with high axial velocity frames (Figure 1 a.). To study this relationship, raster plots of standardized power Doppler time series—referred to as “Carpet plots” (Power, 2017)—are a useful tool (Figure 1 b.). Comparing an example Carpet plot with its corresponding global axial velocity (GV) time series reveals that all power Doppler time series are correlated with the velocity of the brain tissues. Notably, we found that this relationship is non-linear, with high power Doppler intensities consistently associated with high axial velocities (Supplementary Figure 1 a.). This relationship persists even at small velocities and is consistent across all acquisitions in our datasets (Supplementary Figure 1 b.). Importantly, the magnitude of power Doppler surges often exceeds 200% above baseline (Supplementary Figure 1 b.)—far greater than changes observed during typical neural activity—suggesting that these surges cannot be explained by neural activity alone.

To further investigate the temporal characteristics of motion artifacts, we used a finite impulse response (FIR) model to estimate the effect of brain movement on power Doppler intensity (Figure 2 a.). We found that brain movement does not only lead to immediate increases in power Doppler signal, but also has significant effects in directly preceding and following frames. Additionally, we observed a delayed effect, with increases in power Doppler signal peaking at 1.6 seconds in the cortex and hippocampus, and lasting at least 12 seconds in the thalamus.

**Figure 2.**
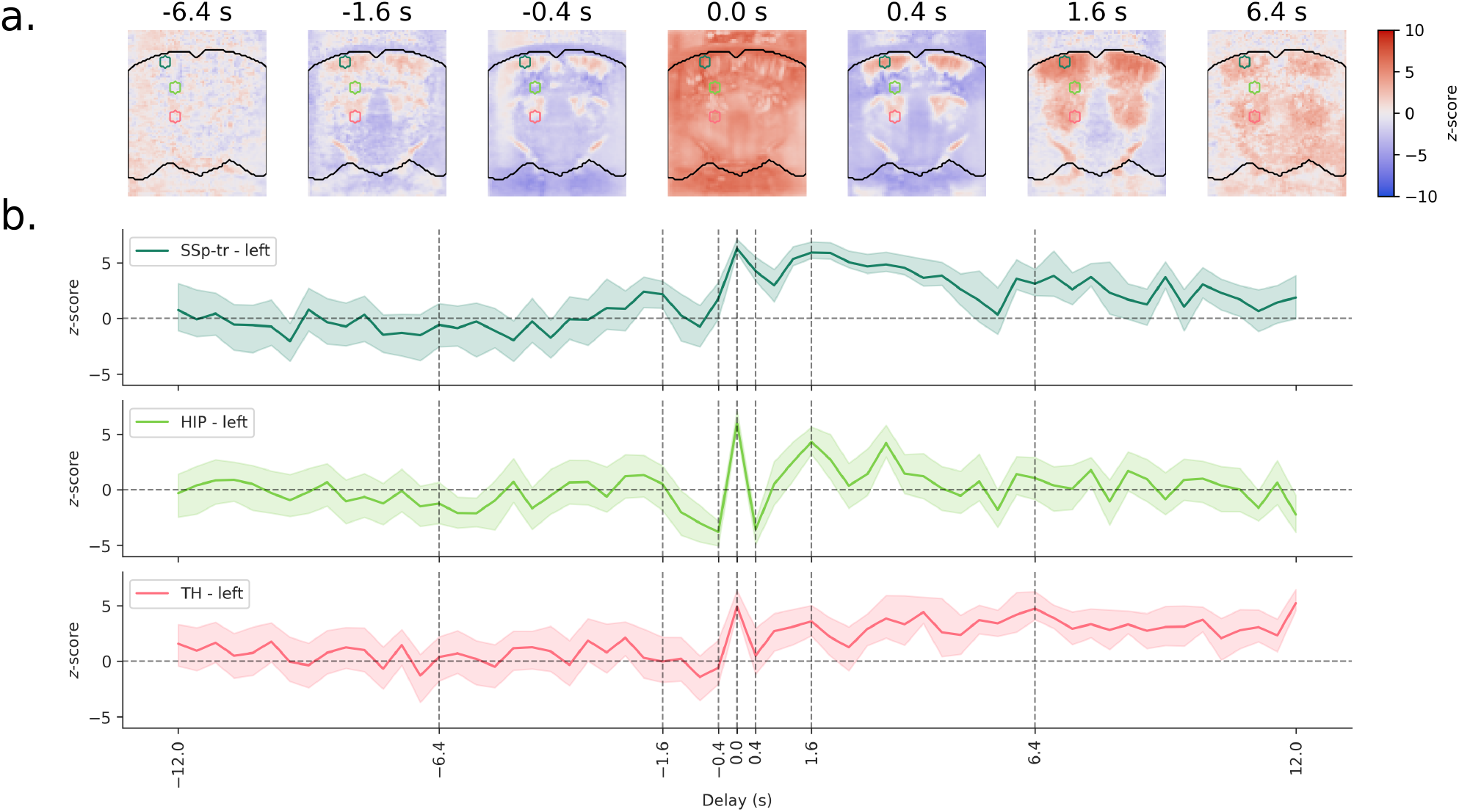
Brain movement induces both immediate and delayed effects on power Doppler signals. (**a**.) A finite impulse response (FIR) model was used to estimate the effect of the brain velocity on power Doppler intensity. Subject-level models were fitted for each mouse of dataset 1, and a group-level model was then fitted across mice. Resulting maps are shown for representative lags (see Supplementary Figure 2 for all lags). At lag 0, brain movement leads to large increases in power Doppler signal across most of the FOV, as seen in Figure 1. However, significant effects are also observed in directly preceding and following frames. A delayed effect is then observed with increases in power Doppler signal across most of the vasculature, peaking at 1.6 seconds in the cortex and hippocampus and lasting up to 12 seconds in the thalamus. (**b**.) Average *z*-score time series are shown for three ROIs in the thalamus, hippocampus, and primary somatosensory cortex, displayed as colored circles in **a**. Shaded areas represent the 95% confidence interval of the mean.

### 3.2 Quality control metrics for artifact detection

To evaluate the impact of motion artifacts in real-world fUSI experiments, quality control (QC) metrics are essential. Here, we propose a set of spatial and temporal QC metrics that may be used to identify acquisitions with high motion artifacts (Figure 3).

**Figure 3.**
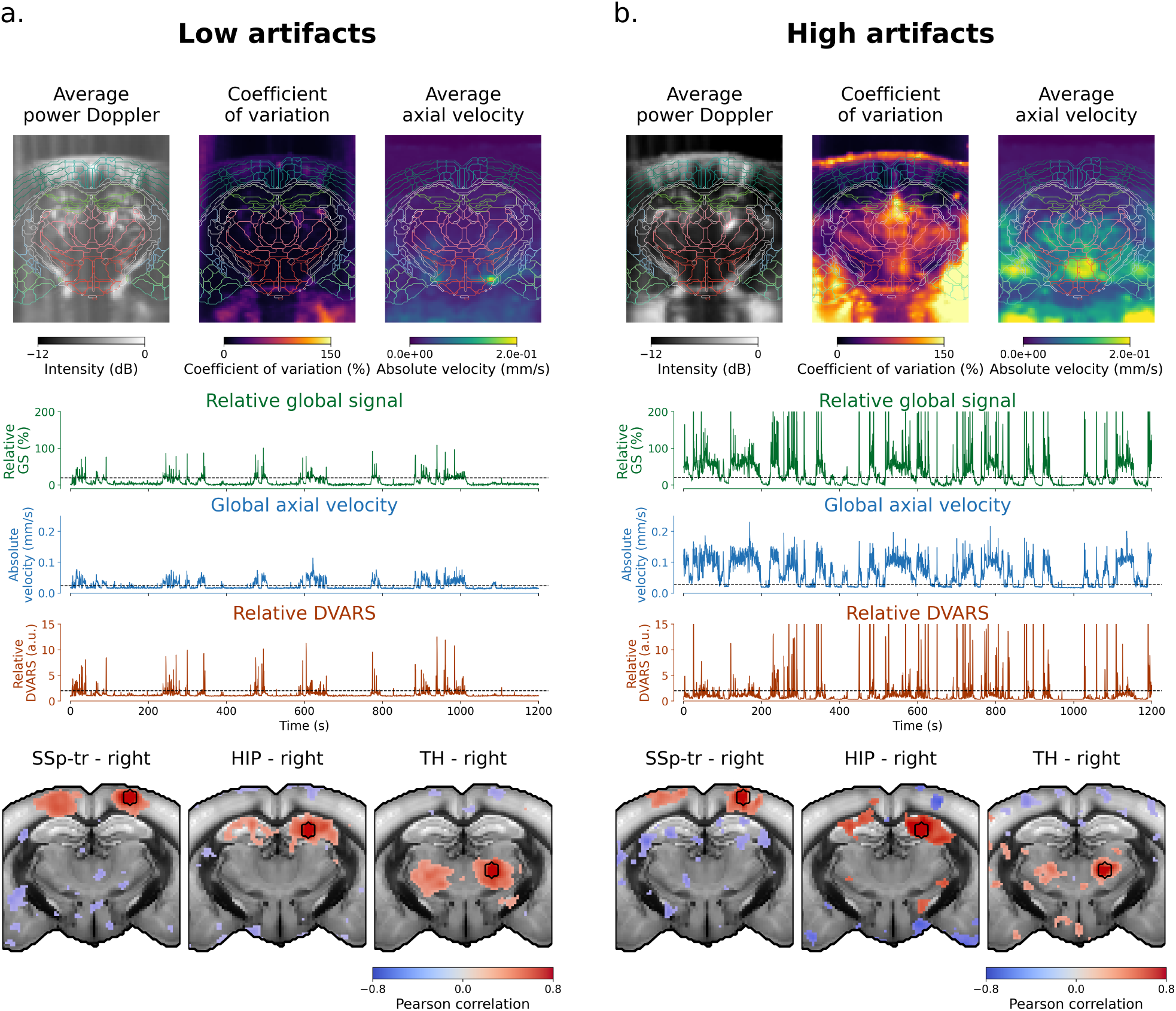
Quality control metrics reveal distinct signatures of motion artifacts. Spatial and temporal QC metrics are displayed for two example acquisitions from dataset 1 with low (**a**.) and high (**b**.) motion artifacts. Both acquisitions were computed using the static SVD60 clutter filter. (**Top**) The top row shows the average power Doppler, CV, and brain axial velocity images. Both acquisitions show loss of power Doppler intensity in lateral and medial parts of the brain, typical of transcranial acquisitions. Both acquisitions also show high CV and high average axial velocity under the brain, where strong flashing artifacts can be observed during movement in power Doppler films. The high motion artifacts acquisition shows higher CV on lateral parts and on the brain edge, and higher axial velocity inside the brain. (**Middle**) The middle row shows the relative global power Doppler signal (rGS), global axial velocity (GV), and relative DVARS (rDVARS) signal. Note the co-occurrence of bursts in all three metrics, indicating that bursts of rGS (and thus rDVARS) are associated with high axial velocities. The high motion artifact acquisition shows more of the bursts, with higher amplitude in average. (**Bottom**) The bottom row shows three example seed-based maps with seeds placed in the primary somatosensory cortex, hippocampus, and thalamus. Seed-based maps were computed using the standard denoising strategy (static SVD60 clutter filtering without brain mask, scrubbing using rGS above 10%, [0.01, 0.1] Hz band-pass filtering, and GSR). Seed-based maps are thresholded to keep 10% of voxels with highest correlation values. Maps are overlaid on the Allen Mouse Brain CCFv3 template. The high motion artifacts acquisition shows loss of bilateral connectivity with more anti-correlated artifacts.

**Figure 4.**
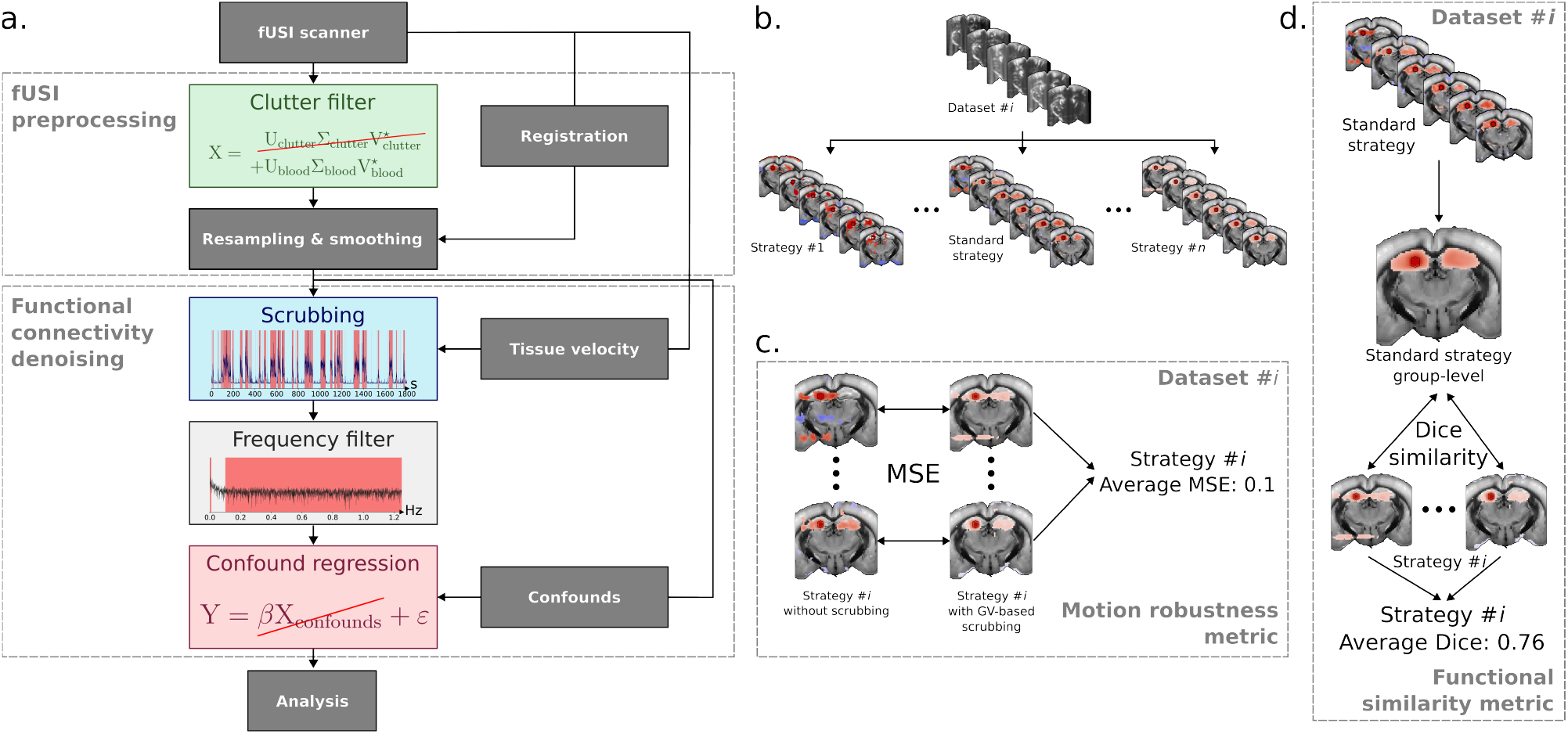
Outline of the denoising strategies and benchmark metrics. (**a**.) General outline of the data denoising strategies benchmarked in this study. (**b**.) Following the general outline in (**a**.), each set of parameters was used to denoise acquisitions from all datasets. Each denoised acquisition was then used to compute 6 seed-based maps. (**c**.) The motion robustness metric was computed for each set of clutter filter, frequency filter, confound regression parameters. The motion robustness metric is the average mean squared error (MSE) across mice between seed-based maps computed with vs. without high-axial-velocity volumes. (**d**.) The FC similarity metric was computed for each strategy and each dataset. The functional similarity metric is the average Dice similarity coefficient across mice between subject-level seed-based maps and the group-level seed-based map computed from the standard denoising strategy.

Spatial QC metrics consist of the average power Doppler intensity—displayed in decibel (dB) scale—, the coefficient of variation, and the average axial velocity. The top row of Figure 3 illustrates these metrics for two example acquisitions from dataset 1 with low and high motion artifacts. Compared to the low artifact example, the high artifact example shows increased coefficient of variation (CV) and axial velocity in large areas of the FOV. On the other hand, the average power Doppler image does not show much difference between the two acquisitions: the high motion artifact acquisition even shows better contrast-to-noise ratio (CNR) compared to the low artifact acquisition. However, the average power Doppler image is useful to identify areas with low signal-to-noise ratio (SNR), such as the lateral and medial parts of the brain. Due to variations in skull thickness, these areas are typically associated with low power Doppler intensity in transcranial acquisitions. Average power Doppler images were therefore used to remove acquisitions with low SNR in ROIs from further analysis (Table 1).

Temporal QC metrics consist of the relative global power Doppler signal (rGS), global axial velocity (GV), and relative DVARS (rDVARS). The middle row of Figure 3 displays these metrics for the same acquisitions. Unsurprisingly, the high motion artifact acquisition exhibits more bursts with higher amplitudes in all three metrics. However, these metrics show different sensitivities to motion artifacts. Notably, surges of rGS are generally on the order of (10 − 60) % with spikes above 200 %, while the GV displays a more consistent increase on the order of 0.1 mm s^−1^. In contrast, the rDVARS strongly penalizes high power Doppler surges, allowing for detection of spikes that are clearly artifactual without removing entire motion periods. Additionally, both GV and rDVARS have stable baselines compared to rGS, making them easier to threshold for motion artifact detection.

Both spatial and temporal QC metrics may be used to identify problematic acquisitions prior to statistical analysis. The bottom row of Figure 3 presents three seed-based maps with seeds placed in the primary somatosensory cortex, hippocampus, and thalamus. Seed-based maps were computed using a standard denoising strategy, similar to those found in recent fUSI studies (see Section 2.9 for details) (Bertolo et al., 2023; Ferrier et al., 2020;

Rabut et al., 2020). The low motion artifact acquisition reveals bilaterally correlated regions with good specificity, whereas the high motion artifact acquisition shows loss of bilateral connectivity and more spurious correlations.

These findings regarding both the temporal and spatial metrics—increased CV and axial velocity in high-artifact acquisitions, variable sensitivity to motion with stable baselines for GV and rDVARS compared to rGS—, as well as the associated degradation in FC specificity, were consistently observed across all four datasets. This highlights the importance of using QC metrics to assess the quality of fUSI data and identify motion-compromised acquisitions.

### 3.3 Outline of the benchmarking pipeline

Next, we designed a benchmarking pipeline to test 792 denoising strategies, created by combining parameters from 3 clutter filters, 3 scrubbing metrics, 2 frequency filtering methods, and 6 confound regression methods (Figure 4 b.). This factorial design allowed us to identify interactions between denoising steps that would not be apparent when testing individual components.

To evaluate their performance for FC analysis, we used two metrics:

▪ First, we created a “motion robustness” metric to quantify the sensitivity of each denoising strategy to motion artifacts. This metric measures the average mean squared error (MSE) between FC maps processed with versus without high-motion volumes. Lower MSE values indicate strategies where removing motion-contaminated volumes makes little difference to the final results, suggesting effective motion artifact suppression through clutter filtering, frequency filtering, and confound regression alone. We used the GV as ground-truth for identifying motion periods, as it directly measures brain movement rather than large signal variations
▪ Second, we developed an FC similarity metric to ensure that motion artifact removal didn’t come at the cost of eliminating neuronally relevant signals. This metric calculates the average Dice coefficient between subject-level FC patterns and group-level FC patterns derived from a commonly used and conservative denoising strategy in the fUSI literature. Higher Dice coefficients indicate better preservation of expected FC patterns.

The combined use of these metrics allowed us to identify denoising strategies that both effectively remove motion artifacts and preserve meaningful signals—a balance that is often challenging to achieve. Additionally, this benchmarking approach provided insights into denoising interactions that would have been impossible to discover through sequential optimization of individual steps.

### 3.4 Robustness to motion artifacts

Our analysis revealed that the motion robustness metric varies significantly across denoising strategies (Figure 5). To facilitate interpretation, we visually selected a threshold of MSE < 0.025 to identify denoising strategies ith satisfactory motion robustness, based on our evaluation of example difference maps. Out of the 216 denoising strategies evaluated, which comprised all possible combinations of parameters from clutter filters, frequency filters, and confound methods, 81 strategies passed this threshold. These successful strategies are represented by filled markers in Figure 5. Notably, the standard denoising strategy, marked by a black star on the figure, exhibited poor robustness to motion, with an average MSE of 0.12.

**Figure 5.**
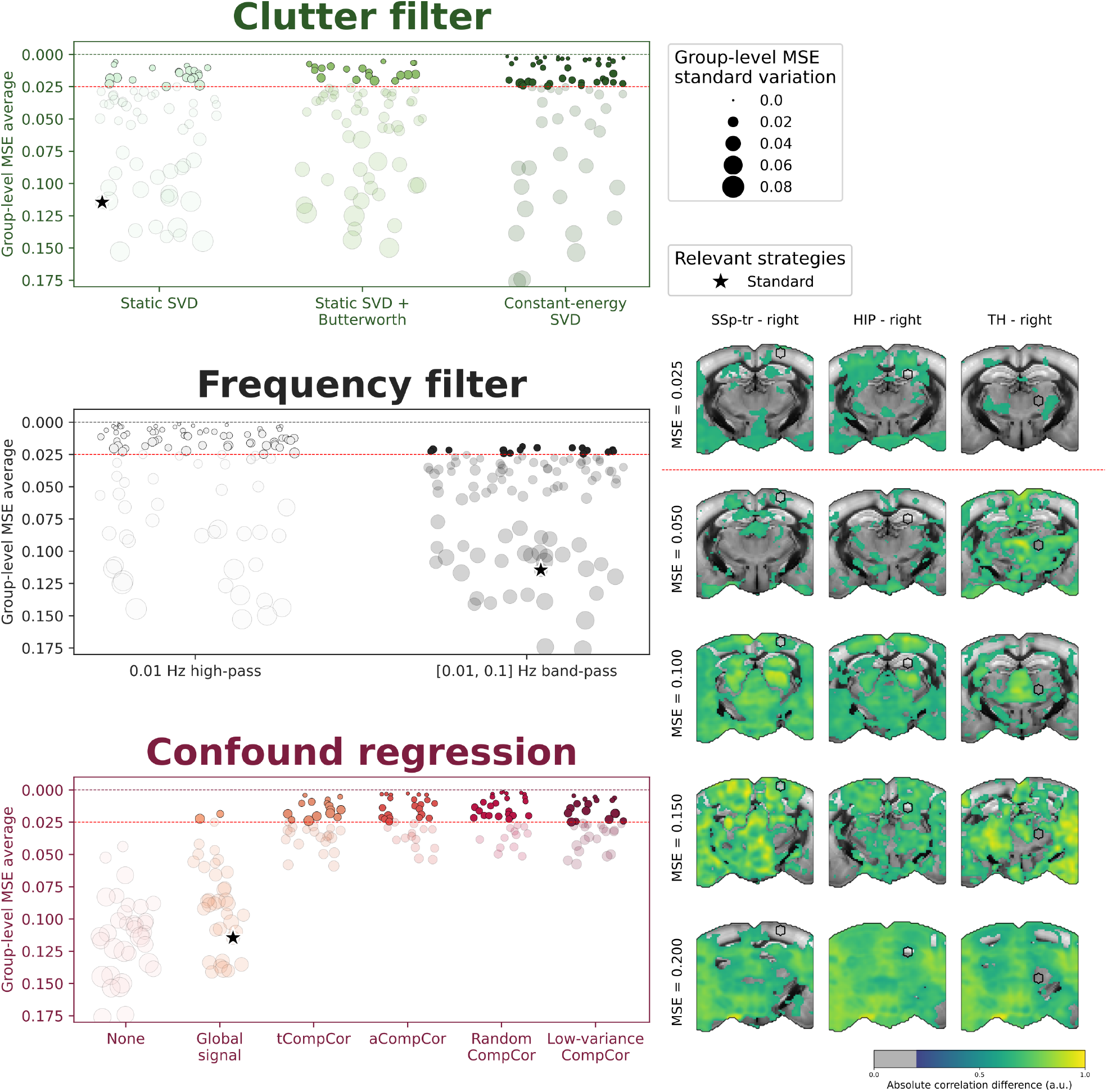
High-pass filtering and CompCor regression methods outperform standard denoising approaches in robustness to motion artifacts. Each denoising strategy was used to compute seed-based maps from all acquisitions of dataset 1. Then, for each set of clutter filter, frequency filter, and confound regression parameters, the average mean squared error (MSE) was computed between subject-level seed-based maps obtained with vs. without high axial velocity volumes (Figure 4 c.). Thus, a parameter set with an MSE of 0 (dotted line) corresponds to a parameter set that will result in similar seed-based maps, regardless of motion artifacts. The radius of the markers corresponds to the group-level MSE standard deviation. All strategies above the MSE = 0.025 threshold (red dotted line) are considered satisfactory and marked by filled markers, and the rest are marked by transparent markers. The standard strategy used in Figure 3 is marked by a black star. Example difference maps are shown on the right for corresponding increments of MSE.

#### 3.4.1 Adaptive clutter filtering and brain masking improves robustness to motion artifacts

Despite the large variability across strategies, we found that all three clutter filter strategies can achieve satisfactory motion robustness. Among the 81 top-performing strategies, 52% used the constant-energy SVD*T* clutter filter, 25% used the static SVD*T* + high-pass clutter filter, and 23% used the static SVD*T* clutter filter. These findings align with suggestions by Baranger and colleagues that adaptive clutter filtering can enhance the robustness of FC analyses against motion artifacts (Baranger et al., 2018).

All three clutter filter thresholds *T* = (20, 40, 60) yielded similar results for the constant-energy SVD*T* clutter filter, with *T* = 40 showing slightly better results overall (Supplementary Figure 3). In contrast, both static clutter filters showed significantly better performance with higher thresholds. This pattern suggests that while lower thresholds may effectively remove clutter components from low-motion volumes, they are insufficient for high-motion volumes. Consequently, static clutter filters require higher thresholds to adequately remove clutter throughout the entire acquisition, which may unfortunately lead to signal loss in low-motion volumes.

Furthermore, our analysis revealed that all satisfactory strategies relying on static SVD clutter filters benefited from computing clutter vectors exclusively from brain voxels, although this improvement is relatively modest (Supplementary Figure 4). On the other hand, 54% of satisfactory strategies relying on the constant-energy SVD*T* filter benefited from a brain mask. Generally, we observed that using a brain mask provides diminishing returns in terms of motion robustness as clutter filtering becomes more stringent. This finding aligns with the assumption that more lenient clutter filtering strategies are susceptible to bias from outliers outside of the brain and may therefore require a brain mask to properly remove clutter from brain voxels. Static SVD clutter filters appear particularly prone to such bias, as the clutter component often exhibits greater variability outside of the brain, especially when the FOV includes adjacent muscles.

#### 3.4.2 High-pass filtering alone outperforms traditional band-pass approaches in the presence of motion artifacts

Among strategies with satisfactory motion robustness, all band-pass filter strategies demonstrated worse robustness to motion artifacts compared to their high-pass filter counterparts (Supplementary Figure 5). This finding suggests that the ringing artifacts and serial correlations introduced by low-pass filtering may be more detrimental in the presence of significant motion artifacts than any potential benefit they might provide in removing high-frequency noise.

#### 3.4.3 CompCor regression substantially outperforms global signal regression

In our analysis of confound regression strategies, all CompCor-based approaches led to substantial improvements in motion robustness compared to global signal regression (GSR) or no-confound regression. Notably, only two GSR strategies managed to pass the MSE < 0.025 threshold, and both relied on the constant-energy SVD*T* clutter filter. This outcome suggests that the global power Doppler signal (GS) does not adequately model motion artifacts. In contrast, we found that just 5 CompCor confounds are sufficient to remove motion artifacts to such an extent that scrubbing becomes unnecessary.

### 3.5 Similarity to group-level FC patterns

Similar to the motion robustness metric, the FC similarity metric exhibits significant variability across denoising strategies (Figure 6). To facilitate interpretation and establish a clear relationship between the motion robustness and FC similarity metrics, we separated the results based on whether scrubbing was used. Additionally, we visually selected a threshold of Dice > 0.75 to identify denoising strategies that produce satisfactory FC patterns. This threshold was determined through examination of example seed-based maps.

**Figure 6.**
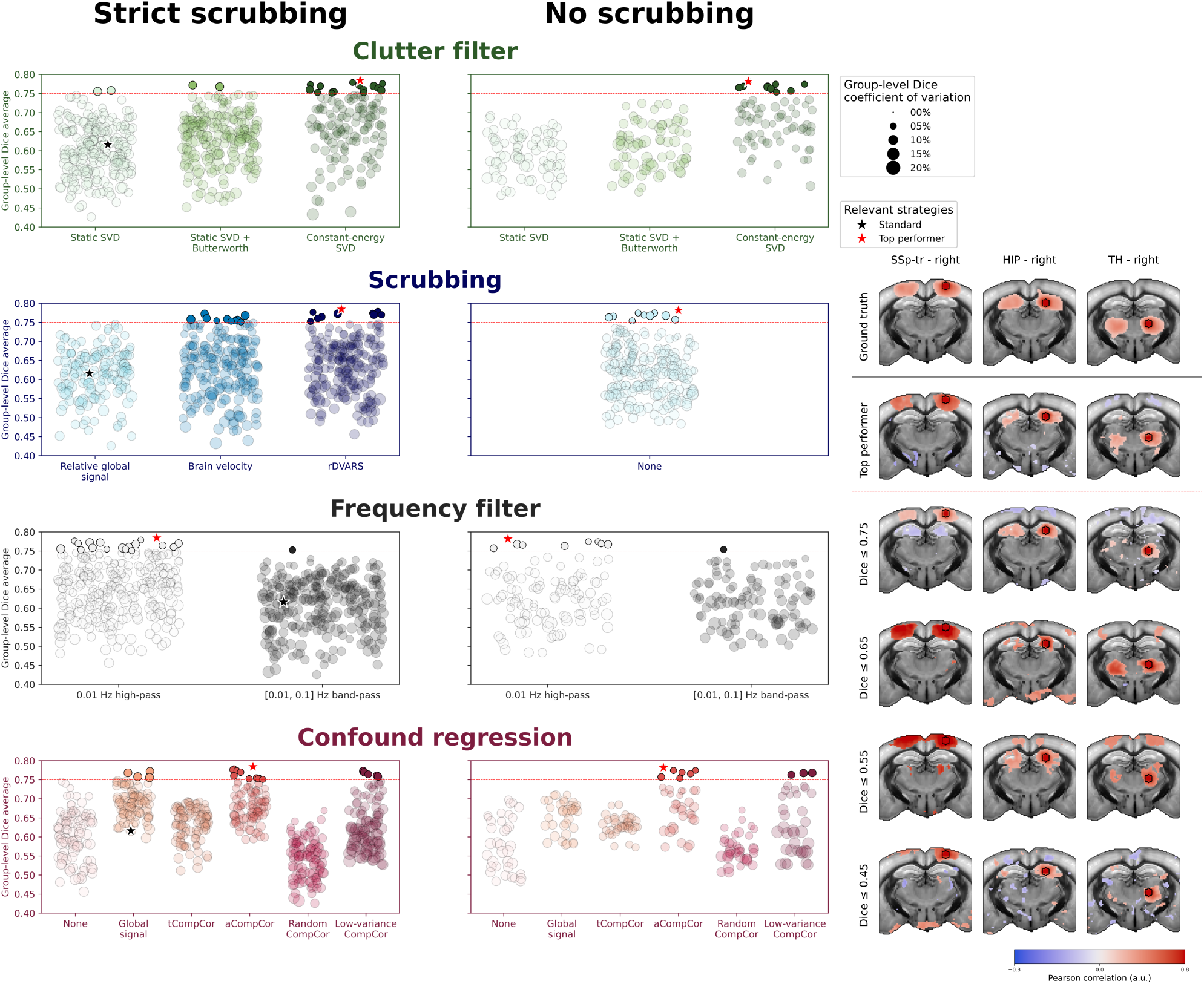
Adaptive clutter filtering, high-pass frequency filtering, and aCompCor and low-variance CompCor regression methods improve FC patterns whether or not scrubbing is used. The FC similarity metric results are separated into strategies with (**left**) or without (**right**) scrubbing. Each set of denoising parameters was used to process acquisitions of dataset 1. For each parameter set, the average Dice similarity coefficient across mice was computed between subject-level seed-based maps and the group-level map obtained with the standard denoising strategy (Figure 4 d.). Thus, a parameter set with a Dice similarity coefficient of 1 corresponds to a strategy that will result in subject-level maps with satisfactory FC regions identical to those in the group-level map. The radius of the markers corresponds to the group-level Dice standard deviation. All strategies above the Dice = 0.7 (red dotted line) threshold are marked by filled markers, and the rest are marked by transparent markers. The standard strategy used in Figure 3 is marked by a black star, and respective top performers for the “strict scrubbing” and “no scrubbing” cases are marked by a red star. Example seed-based maps are shown on the right for corresponding increments of Dice similarity coefficient, along with the group-level ground-truth.

Out of these 792 denoising strategies, only 30 demonstrated satisfactory FC patterns, which are indicated by filled markers on Figure 5. The standard denoising strategy, marked by a black star on Figure 6, falls below our threshold with a Dice coefficient of approximately 0.62. Interestingly, the top-performing strategy for the FC similarity metric employs the same parameters regardless of whether scrubbing is used: constant-energy SVD*T* clutter filter with brain mask, high-pass filter, and aCompCor confound regression.

#### 3.5.1 Adaptive filters maintain connectivity networks without scrubbing

Compared to the motion robustness metric, the FC similarity metric differentiates more clearly between clutter filter methods. Here, the constant-energy SVD*T* clutter filter represents 87% of the top performers, clearly outperforming static clutter filters. In contrast, only 2 static SVD*T* and 2 static SVD*T* + high-pass clutter filter strategies demonstrate satisfactory FC patterns when scrubbing is used, while no static clutter filter achieves acceptable performance without scrubbing. When considered alongside the motion robustness results, these findings further indicate that static clutter filters may be poor choices for FC analyses in cases where large variations in clutter components are present.

Moreover, the few static clutter filters that do achieve satisfactory FC patterns benefit from higher thresholds similarly to what we observed using the motion robustness metric. Meanwhile, the constant-energy SVD*T* clutter filter demonstrated similar performance with thresholds of *T* = 20 and *T* = 40, but showed a decline in FC similarity at *T* = 60. These observations strengthen our assumption that static clutter filters may remove relevant signal from low-motion volumes due to their reliance on high thresholds for effective clutter removal across full acquisitions.

Finally, all static SVD-based and 83% of constant-energy SVD-based strategies that produce satisfactory FC patterns benefit from using a brain mask to compute clutter vectors (Supplementary Figure 4). Although this benefit remains modest for the best-performing strategies, this result aligns with the trend observed in the motion robustness metric.

#### 3.5.2 relative DVARS and global axial velocity metrics provide more effective scrubbing

Our analysis revealed that GV, rDVARS, and no-scrubbing strategies can lead to satisfactory FC patterns, each representing 33% of top-performing strategies. The fact that no top-performing strategy used rGS-based scrubbing is particularly noteworthy, as it suggests that rGS may not be suitable for FC analyses.

Interestingly, scrubbing appears to be optional for strategies that employ the constant-energy SVD*T* clutter filter. Top performers for both scrubbing and no-scrubbing scenarios use the constant-energy SVD*T* clutter filter with nearly identical results. This supports our previous observations that adaptive clutter filters can effectively mitigate motion artifacts upstream, resulting in very few volumes requiring removal.

In contrast, static clutter filters demonstrate a clear benefit from scrubbing, with no top-performing strategy using static clutter filters without scrubbing. This suggests that static clutter filters may be insufficient for removing motion artifacts on their own, necessitating scrubbing to eliminate residual artifacts.

#### 3.5.3 Low-pass filtering degrades connectivity in all configurations

All best-performing strategies benefit from using high-pass filtering compared to band-pass filtering, confirming our observations from the motion robustness metric regarding the detrimental effects of low-pass filtering in the presence of motion artifacts.

#### 3.5.4 aCompCor and low-variance CompCor consistently preserve FC patterns

When scrubbing is used, only global signal regression (GSR), aCompCor, and low-variance CompCor strategies produce satisfactory FC patterns. Surprisingly, other CompCor strategies yield poor results, despite their good performance in terms of motion robustness. In particular, random CompCor performs worse than strategies that do not regress any confounds at all. This indicates that both methods remove excessive signal of interest, resulting in an almost complete loss of correlated regions in the seed-based maps.

When scrubbing is not used, the performance of GSR strategies decreases significantly, while aCompCor and low-variance CompCor strategies continue to yield comparably good results. Overall, 60% of the best-performing strategies employ aCompCor confound regression, regardless of whether scrubbing is used. This suggests that aCompCor represents a robust confound regression method for FC analyses in awake mice, effectively balancing the removal of motion artifacts with the preservation of signals of interest.

Notably, low-variance CompCor proved useful only when combined with the constant-energy SVD*T* clutter filter. This may indicate that areas of low temporal variance do not contain sufficient information to properly model strong motion artifacts, yet are adequate for removing other noise sources when used in conjunction with adaptive clutter filters.

### 3.6 Benchmarking summary

To provide a comprehensive overview of denoising strategy performance, we summarized the motion robustness and FC similarity metrics into a single scatter plot (Figure 7). Our analysis revealed that only 26 out of the 792 benchmarked strategies demonstrated both satisfactory motion robustness and FC similarity. These top-performing strategies are highlighted by filled markers in the inset axes of Figure 7.

**Figure 7.**
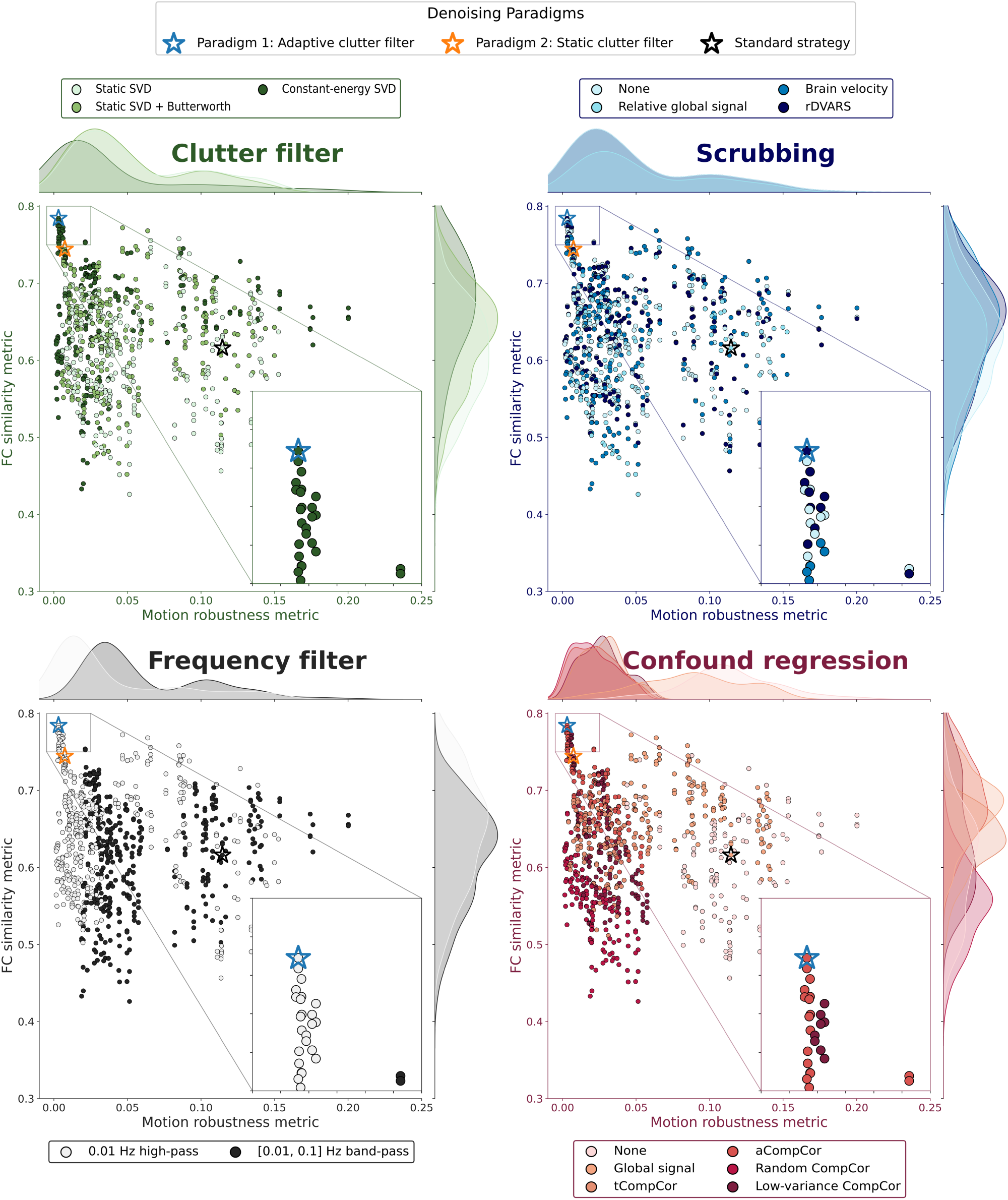
Only 26 of 792 denoising strategies achieve both satisfactory motion robustness and FC pattern preservation, all employing adaptive SVD clutter filtering. Metric values from Figure 5 and Figure 6 are summarized into a single scatter plot. The same plot is shown four times, with color corresponding to the different denoising steps. The inset axes shows strategies that pass the thresholds for both metrics. The four global strategies and the standard strategy are marked by stars.

Examining top-performing strategies more closely, we found that the constant-energy SVD *T* clutter filter predominated, being used in all 26 strategies. All three thresholds were represented among these successful strategies, with *T* = 20 accounting for 42%, *T* = 40 for 35%, and *T* = 60 for the remaining 23%.

Further analysis revealed that all scrubbing approaches but rGS were represented: 39% used rDVARS-based scrubbing, 23% used GV-based scrubbing, and 38% employed no scrubbing at all.

Regardless of the scrubbing approach, all best-performing strategies consistently employed high-pass filtering and either aCompCor or low-variance confound regression methods.

### 3.7 Optimized denoising paradigms perform consistently across multiple datasets

Having identified the key parameters that influence denoising performance, we sought to develop practical recommendations for fUSI data denoising. For this purpose, we categorized strategies into two general paradigms, based on their use of adaptive or static clutter filtering. Indeed, we expect the choice of clutter filter to be the most controversial among the benchmarked denoising parameters, primarily because the lack or large size of beamformed I/Q data or the need for non-normalized CBV time series (for example, to retain global variations) may restrict the use of adaptive filters. Then, we selected best-performing strategies for each paradigm, which we refer to as paradigms 1 and 2 for adaptive and static clutter filtering, respectively. Parameters for the proposed paradigms, along with the standard denoising strategy, are summarized in Table 2.

**Table 2:** Parameter configurations for the two optimal denoising paradigms. Each paradigm represents an optimized strategy for denoising fUSI data based on the motion robustness and FC similarity metrics. The parameters of the standard strategy are also included for comparison. Paradigm 1 employs adaptive clutter filtering while paradigm 2 uses static clutter filtering. Since clutter filtering impacts the relative global power Doppler signal (rGS) and relative DVARS (rDVARS) scrubbing metrics, the across-mice average and standard deviation of the temporal degrees of freedom (tDOF) are shown each strategy. Note that both paradigm 1 and 2 converged on the same parameters regardless of the clutter filter.

|  | Clutter filter | Scrubbing | Frequency filter | Confound regression | tDOF | Motion robustness | FC similarity |
| --- | --- | --- | --- | --- | --- | --- | --- |
| Standard strategy | Static SVD60 | rGS | (0.01 – 0.1) Hz band-pass | GS | $(52 \pm 8) \%$ | $1.2 \cdot 10^{-1}$ | 0.62 |
| Paradigm 2 | Static SVD60 + high-pass + brain mask | rDVARs | 0.01 Hz high-pass | aCompCor | $(91 \pm 4) \%$ | $7.4 \cdot 10^{-3}$ | 0.74 |
| Paradigm 1 | Constant-energy SVD40 + brain mask | rDVARs | 0.01 Hz high-pass | aCompCor | $(95 \pm 3) \%$ | $3.1 \cdot 10^{-3}$ | 0.78 |

Our analysis revealed that the best-performing strategy for paradigm 1 relies on the constant-energy SVD40 clutter filter with brain mask, while paradigm 2 employs the static SVD60 clutter filter with high-pass filtering and brain mask. Beyond these differences in clutter filtering, both optimal strategies share common parameters: they employ high-pass filtering, rDVARS-based scrubbing, and aCompCor confound regression. This consistency in optimal parameters across different paradigms underscores their effectiveness fUSI data denoising.

A notable finding is that both paradigms retain significantly more time points (paradigm 1: (95 ± 3) % paradigm 2: (91 ± 4) %) compared to the standard strategy ((52 ± 8) %). This increase in temporal degrees of freedom is particularly important for experimental designs that rely on short acquisitions, but also to reduce the required duration of regular awake paradigms.

Moreover, removing scrubbing entirely from paradigm 1 does not significantly affect the performance, resulting in almost identical FC similarity on average. Thus, for experimental conditions where scrubbing may introduce bias, such as behavioral experiments, scrubbing can be omitted without compromising the results. In contrast, removing scrubbing from paradigm 2 leads to a significant drop in FC similarity, which is likely due to the static clutter filter’s inability to remove motion artifacts effectively.

To assess the generalizability of these paradigms, we validated their performance by computing the motion robustness and FC similarity metrics on datasets 2 and 3 (Figure 8). The results demonstrated consistent performance patterns across all three datasets. Importantly, paradigm 1 surpassed the thresholds for both metrics across all datasets, confirming its robustness. In contrast, paradigm 2 exhibited poorer performance and crossed the threshold only for dataset 2.

**Figure 8.**
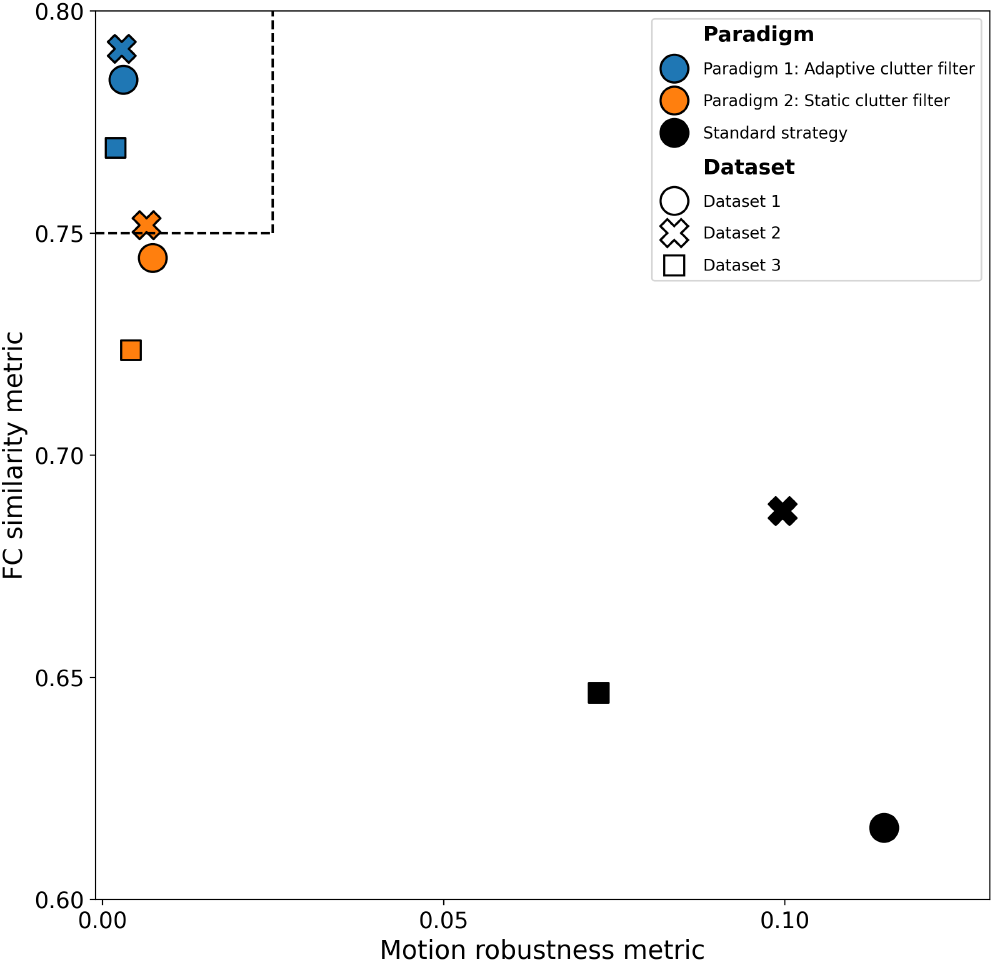
Optimized denoising paradigms demonstrate consistent performance improvements across multiple datasets compared to standard approaches. The two general denoising paradigms and the standard strategy are shown on a scatter plot of FC similarity vs. motion robustness. Strategies are indicated by marker color, while datasets are indicated by marker shape. The respective metric thresholds are shown as dashed lines.

Overall, both optimized paradigms substantially reduced cross-dataset variability compared to the standard strategy, which consistently performed poorly across all datasets. This validation confirms that our methodological recommendations are generalizable and provide more reliable results than currently established strategies.

### 3.8 Optimized denoising paradigms reveal networks obscured by artifacts in standard approaches

To illustrate the practical implications of the different denoising strategies on FC analyses, we divided datasets 1, 2, and 3 into two groups based on consisting of the 30% most, respectively least, motion contaminated acquisitions. Motion contamination was determined by the average GV metric and spatial QC metrics (Figure 3). We then computed group-level seed-based maps for the standard denoising strategy (with and without scrubbing) and the optimized paradigms 1 and 2. Seeds were placed in three ROIs: the primary somatosensory cortex, hippocampus, and thalamus. This approach allowed us to assess the impact of motion artifacts on the resulting FC maps (Figure 9). Additionally, we computed group-level seed-based maps for dataset 4 using the standard denoising strategy and the optimized paradigms 1 and 2 (Figure 10). Given the low number of mice in this dataset and the fact that the standard strategy leads to results too artifactual to be used as ground-truth, dataset 4 was not included in the benchmark. Instead, dataset 4 was used to illustrate the performance of the optimized paradigms on 3D fUSI data.

**Figure 9.**
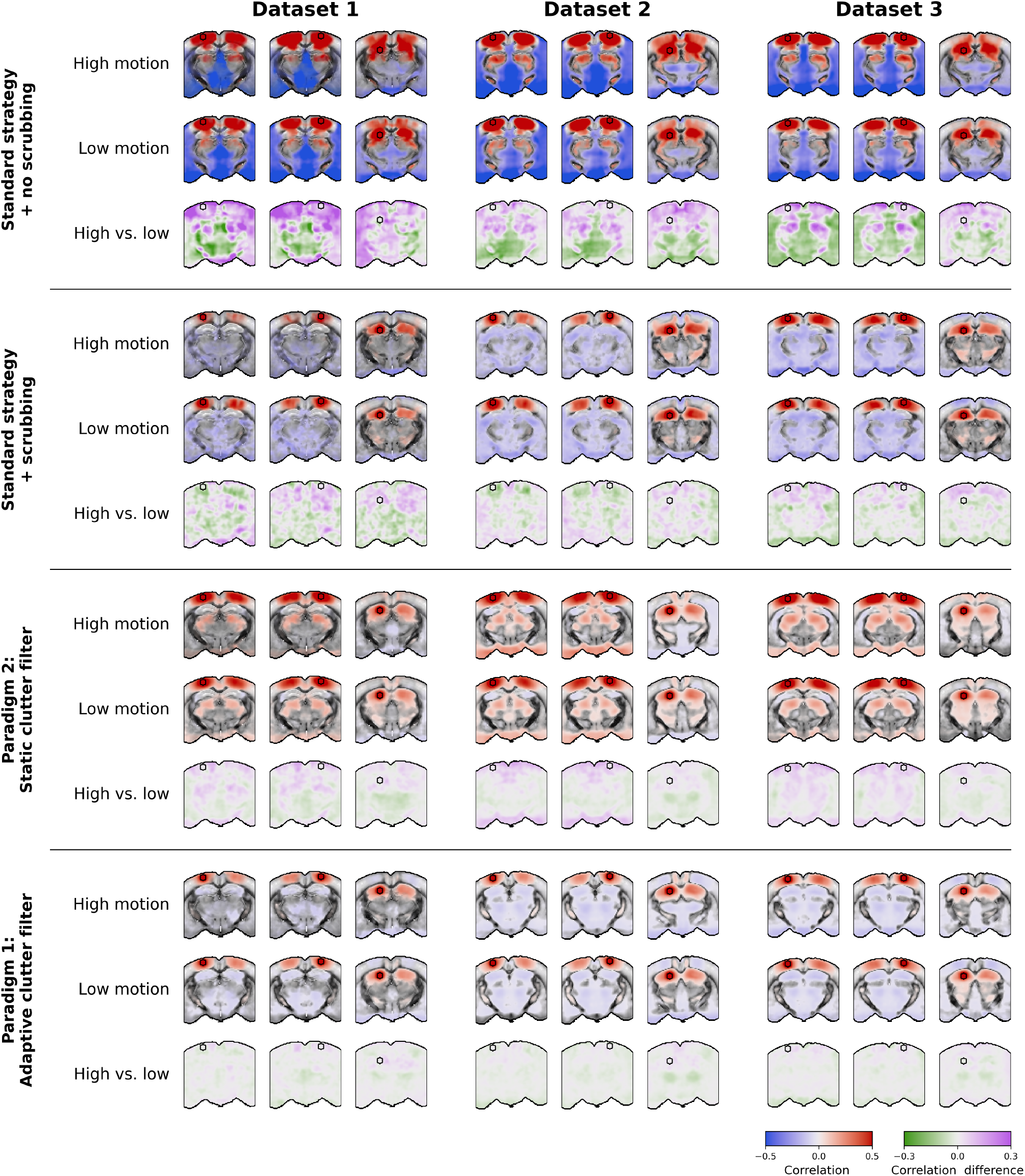
Optimized denoising paradigms preserve bilateral connectivity patterns and reduce artifactual anti-correlations in 2D seed-based maps compared to the standard strategy. Group-level seed-based maps are shown for the primary somatosensory cortex, hippocampus, and thalamus. Group-level maps were computed by averaging correlations (with Fisher transformation) within each mouse. Transparency thresholding was then applied, keeping significant values opaque while adjusting the opacity of non-significant values relatively to their *p*-value (*t*-test on Fisher-transformed correlations, family-wise error rate (FWER) controlled at 5%). Difference maps between high- and low-motion groups are computed by subtracting the average seed-based maps of the high-motion group from the low-motion group.

**Figure 10.**
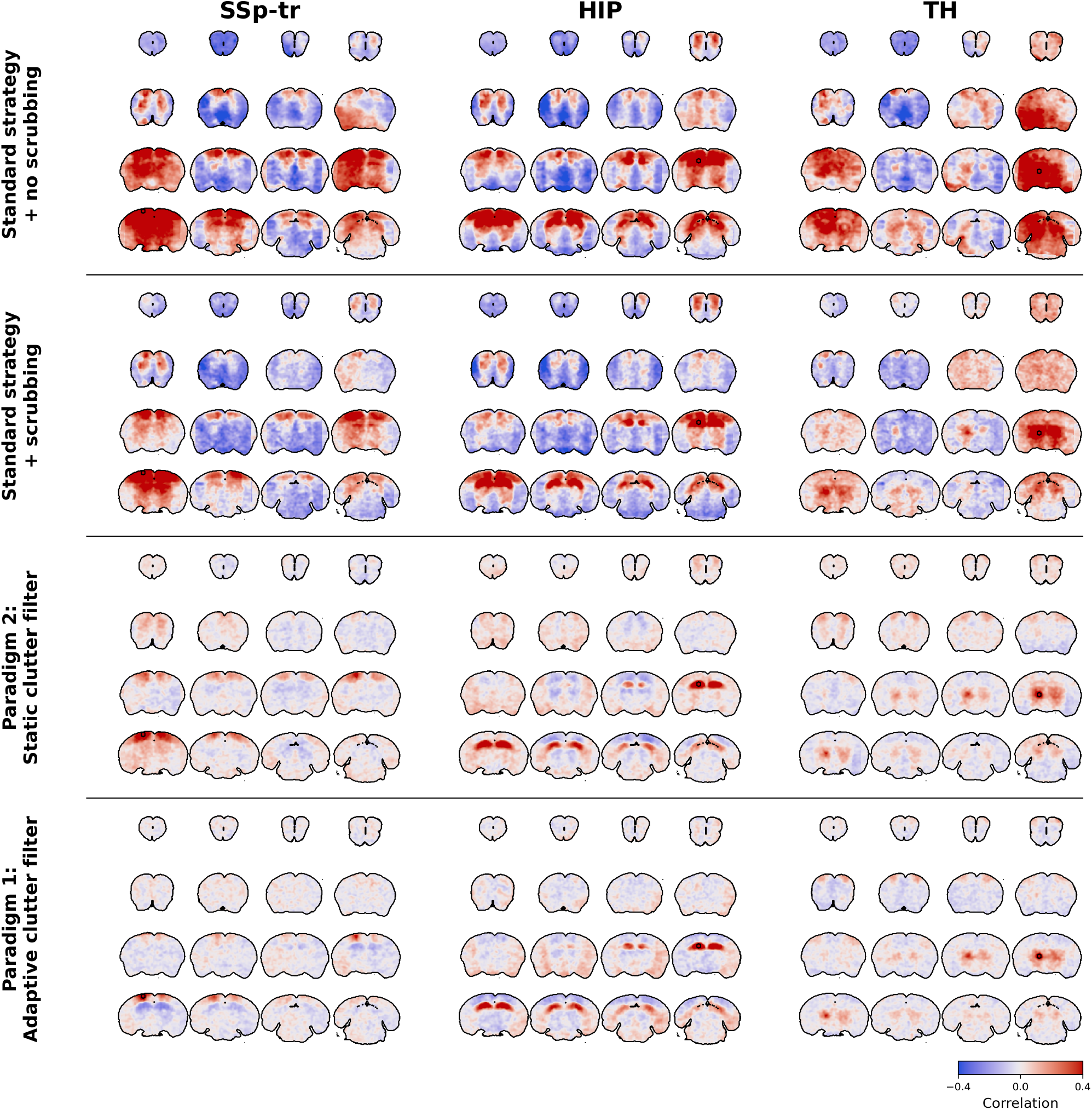
Optimized denoising paradigms enable recovery of anatomically plausible functional networks in 3D acquisitions where the standard strategy fails drastically. Group-level seed-based maps are shown for the primary somatosensory cortex, hippocampus, and thalamus. Group-level maps were computed by averaging correlations (with Fisher transformation) within each mouse.

As shown in Figure 9, the standard denoising strategy exhibited the largest differences in FC patterns between high- and low-motion groups. When not using scrubbing, the standard strategy produced extreme positive and negative correlations across very large areas of the brain, with clearly artifactual patterns between groups. When using scrubbing, the standard strategy produced plausible patterns that constituted the ground-truth of this study. However, strict scrubbing was necessary, as around half of the time points were removed from the analysis on average Table 3. Additionally, reduced bilateral correlations can be observed in some cortical areas from the high-motion groups in datasets 1 and 2.

**Table 3:** Distribution of acquisition and mice among motion groups for datasets 1, 2, and 3.

| Dataset | Motion group | Number of mice | Number of acquisitions |
| --- | --- | --- | --- |
| Dataset 1 | Low | 16 | 35 |
|  | High | 10 | 35 |
| Dataset 2 | Low | 30 | 39 |
|  | High | 35 | 39 |
| Dataset 3 | Low | 32 | 95 |
|  | High | 32 | 95 |

**Table 4:**
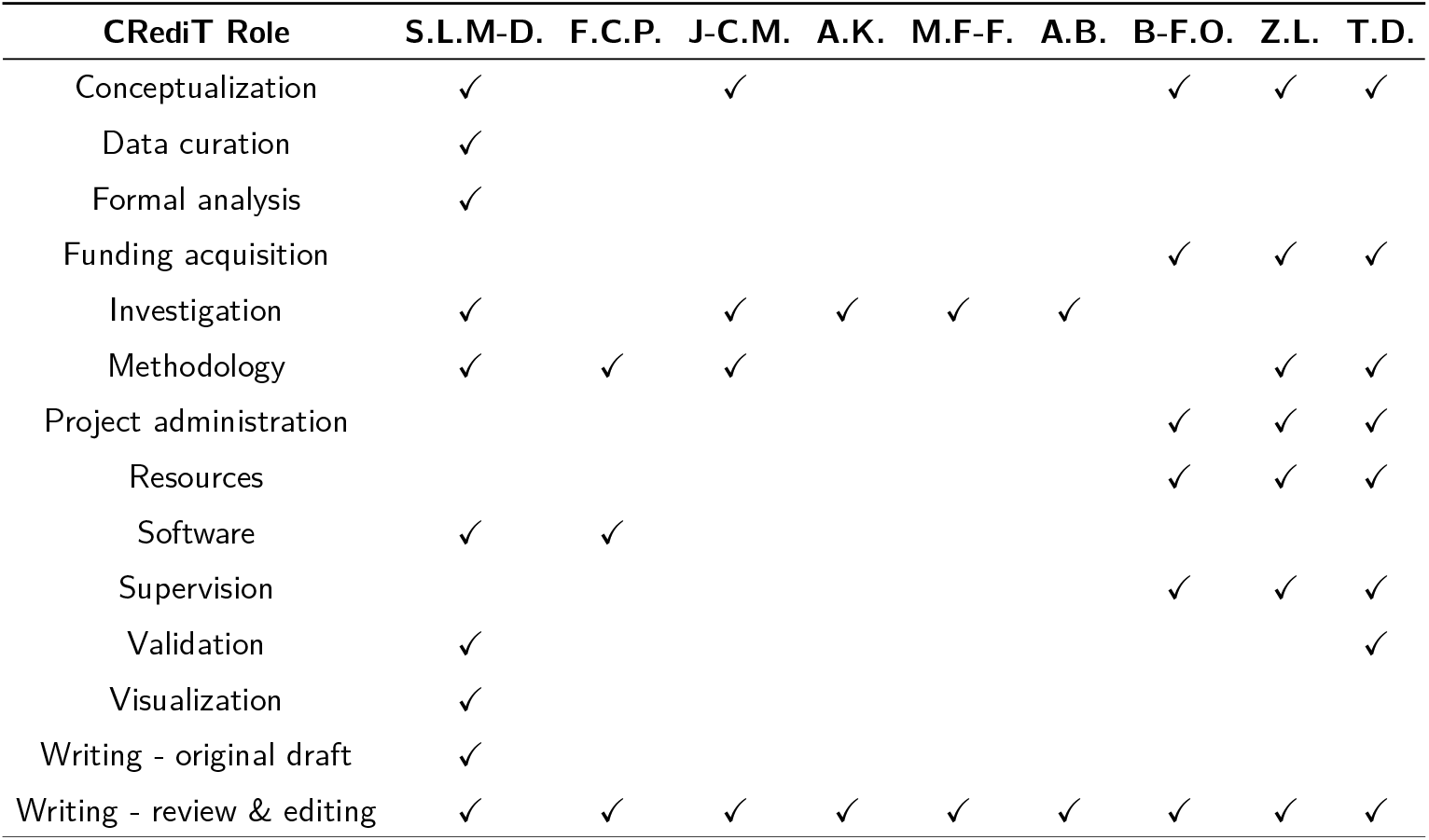
Author Contributions. Author contributions according to CRediT (Contributor Roles Taxonomy). S.L.M-D: Samuel Le Meur-Diebolt; F.C.P: Felipe Cybis Pereira; J-C.M: Jean-Charles Mariani; A.K: Andrea Kliewer; M.F-F: Miguel Farinha-Ferreira; A.B: Adrien Bertolo; B-F.O: Bruno-Félix Osmanski; Z.L: Zsolt Lenkei; T.D: Thomas Deffieux.

Moreover, many anti-correlated regions are observed, in particular in ventral regions, which are likely due to flashing artifacts from lower cerebral veins leaking into the brain.

In dataset 4, the standard strategy was unable to remove motion artifacts whether or not scrubbing was used. The resulting seed-based maps were highly artifactual, with large correlations between the brain slice containing each seed region and brain slices corresponding to the same sub-array of the multiarray probe.

In contrast, paradigm 2 produced lower differences between high- and low-motion groups and less anti-correlated regions. Moreover, paradigm 2 revealed thalamo-cortical correlations in all datasets that were absent when using the standard strategy. However, correlated regions seem overall less specific in datasets 1 to 3, and spurious correlations in ventral regions are still visible, in particular in dataset 2.

Finally, paradigm 1 produced the most reliable results, with bilateral patterns resembling those obtained using the standard strategy, but with more specificity in both positive and negative correlations, and lower differences between high- and low-motion groups. Note however that anti-correlated patterns should be interpreted with caution, as they may be due to the constant-energy SVD*T* clutter filter. Indeed, by normalizing the power Doppler signal, the constant-energy SVD*T* clutter filter might introduce negative correlations in a similar fashion to GSR (Murphy & Fox, 2017).

Interestingly, some differences between high- and low-motion groups seem to be physiological, characterized by increased thalamo-cortical correlations and decreased thalamo-hippocampal correlations in the high-motion group. This pattern is consistent is particularly visible in datasets 2 and 3 when using both optimized paradigms and could be related to neural activity changes during motion. This finding suggests that paradigms 1 and 2 are rather able to efficiently remove motion artifacts while preserving relevant neurophysiological signals even during motion periods.

Overall, these visual comparisons align with our quantitative metrics, confirming that the optimized paradigms may produce more anatomically plausible and statistically robust FC maps while removing very few time points. Furthermore, these results demonstrate that appropriate denoising not only improves statistical metrics but also enhances the biological interpretability of fUSI data, allowing more accurate identification of functional networks in the awake mouse brain.

## 4 Discussion

The successful implementation of functional ultrasound imaging (fUSI) in awake rodents has been fundamentally limited by motion artifacts. These artifacts manifest as intense, synchronized bursts in the power Doppler signal, inducing systematic biases that compromise both individual subject analyses and group-level comparisons. Our study reveals insights into the spatial and temporal characteristics of motion artifacts in awake transcranial fUSI, providing a foundation for their successful mitigation. Importantly, our benchmark of 792 denoising strategies indicates that motion artifacts can be significantly mitigated by improved clutter filtering, frequency filtering, and confound regression.

### 4.1 Motion artifacts stem from complex interactions beyond simple tissue movement

A better understanding of motion artifacts in awake fUSI data is crucial to develop tailored denoising strategies. Our results indicate that brain motion induces large, simultaneous, but spatially non-uniform increases in both axial velocity and power Doppler intensity. Increases in axial velocity are observed across the whole brain tissue, but particularly prominent in ventral areas, likely reflecting the head-fixed setup which constrains motion from the dorsal part the skull. This pattern suggests that current head-fixation approaches, while preventing gross movements, still permit subtle deformations of brain tissue.

In contrast, surges of relative power Doppler signal above 100% are mainly observed near the skull (both dorsal and ventral), in large veins, and in lateral areas closer to muscles. While the temporal relationship between brain motion and power Doppler intensity surges is clear, the discrepancy between their spatial patterns suggests that motion artifacts in power Doppler signals are not solely related to tissue movements. Rather, they likely arise from complex interactions between tissue motion, neuronal activity, acoustic reflections, and the signal processing pipeline—especially clutter filtering. This multi-factorial nature calls for further investigation employing another ground-truth measure of CBV as intrinsic optical signal imaging (IOSI) or laser Doppler flowmetry (LDF) (Takuwa et al., 2014)—to disentangle the contributions of motion artifacts and neurovascular coupling.

Finally, we report a delayed effect of animal motion, manifested as power Doppler changes persisting for several seconds after the movement itself. This observation has important methodological implications. First, we found that motion artifacts are characterized by an immediate increase in power Doppler signal, coinciding with the onset of axial velocity surges. This is what we refer to as “motion artifacts”. However, we also observed a delayed increase in power Doppler signal across several brain regions, with longer persistence in cortical and thalamic regions. Notably, the observed 1.6 s delay suggests a neuronal origin, which is consistent with previously reported hemodynamic response function (HRF) delays measured using fUSI in mice (Nunez-Elizalde et al., 2022).

### 4.2 Quality control metrics are essential for reliable assessment of fUSI data

A critical aspect of establishing fUSI as a reliable neuroimaging technique for awake animals is the development of appropriate quality control (QC) metrics. While such metrics are well-established in fMRI (Parkes et al., 2018; Power et al., 2014), fUSI has yet to receive similar attention. Our study provides a first set of simple and interpretable QC metrics that can be used to assess the quality of fUSI acquisitions.

#### 4.2.1 Coefficient of variation and average velocity images are more informative than average power Doppler images

Our analysis reveals several counterintuitive properties of spatial QC metrics in fUSI data. Although the average power Doppler image is generally the first spatial metric to be inspected when processing an acquisition, we found that it fails to effectively discriminate acquisitions with strong motion artifacts. This limitation stems both from the absence of visible brain motion and from the sporadic nature of motion artifacts, where transient signal surges tend to average out over time. However, visual inspection of average power Doppler images still serves an important purpose in identifying acquisitions with low SNR, for example due to variations in skull thickness. Indeed, such acquisitions contribute significantly to spurious group-level variability and should be excluded through careful visual inspection.

Similarly, the standard deviation of power Doppler intensity and traditional temporal signal-to-noise ratio (tSNR) metrics—widely used in other neuroimaging modalities—prove problematic for fUSI QC. The intrinsic positive correlation between temporal average and standard deviation of power Doppler signals creates a paradoxical situation where areas with low signal—such a as “shadow zones” due to skull attenuation, or the ultrasound gel— exhibit high temporal signal-to-noise ratio (tSNR) values, while the vasculature itself shows low tSNR. This relationship fundamentally differs from other neuroimaging modalities like fMRI, where higher tSNR generally indicates better signal quality (Murphy et al., 2007; Welvaert & Rosseel, 2013).

This correlation between signal magnitude and variance can be understood through the physical principles of power Doppler imaging, where the signal is proportional to both the number of scatterers in the voxel and the ultrasound amplitude. Consequently, areas with high power Doppler intensity (corresponding to large blood vessels) naturally exhibit greater physiological variations. This relationship represents a fundamental property of the technique rather than a quality issue, which must be considered when analyzing fUSI data.

Our observations suggest that the temporal CV as signal standard deviation over mean, effectively the inverse of the tSNR—offers a more informative metric for identifying motion artifacts. Indeed, the CV compares signal variance—which is very high in flashing artifacts —to the signal mean—which is never zero, even low-intensity areas. Thus, the CV provides an interpretable measure that can be compared against known physiological variations in CBV, making it particularly valuable for identifying regions with abnormal signal variations. Combined with the average axial velocity image—which provides a ground-truth measure of tissue movement, akin to the framewise displacement (FD) in fMRI—these metrics form a complementary set for spatial QC, allowing diagnosis of clutter artifacts, acoustic reflections, or problems with head fixation.

#### 4.2.2 Velocity-based metrics provide more reliable movement detection

For temporal QC assessment, we found that the relative global power Doppler signal (rGS), global axial velocity (GV), and relative DVARS (rDVARS) metrics are generally highly correlated in awake task-free experiments. However, these metrics differ significantly in their utility across experimental contexts.

While rGS effectively identifies motion artifacts in task-free conditions by comparing with known physiological CBV variations, it becomes problematic in experimental paradigms where legitimate CBV changes can occur independently of motion, such as during sensory stimulation, pharmacological manipulation, or rapid eye movement (REM) sleep.

In contrast, the GV metric is a particularly valuable ground-truth measure for identifying motion-contaminated volumes, with a notably stable baseline that facilitates straightforward thresholding. Although theoretically susceptible to physiological signals like cardiac pulsatility and respiration (Zucker et al., 2025), our analysis suggests that GV is predominantly driven by brain movements. Moreover, the absolute averaging approach used in computing this metric—leading to the sample sampling frequency as power Doppler acquisitions—effectively acts as a low-pass filter that attenuates higher-frequency physiological oscillations.

Implementing GV-based QC across different experimental setups presents challenges that require careful consideration. Notably, brain tissues exhibit region-dependent movement amplitudes, necessitating proper standardization when comparing GV across different FOVs. Additionally, the GV is further limited by the fact that only the *z*-axis component of velocity is measured, and by the susceptibility of the Kasai autocorrelation estimator to noise. To mitigate these limitations, we recommend filtering beamformed I/Q signals prior to computing axial velocities to avoid biases from high-frequency noise (for example, electrical noise).

The rDVARS metric provides another valuable approach for detecting signal surges, with properties complementary to GV. rDVARS is a standardized version of DVARS, proposed by Nichols and colleagues (Nichols, 2017), which we found sufficient for cross-acquisition comparisons and identification of motion artifacts in our datasets. Like GV, rDVARS is easy to threshold thanks to its stable baseline. However, rDVARS is more sensitive to large spikes in power Doppler signals, due to its definition as the RMS of the temporal derivative of the signal intensity. Thus, rDVARS may be particularly useful in detecting sudden power Doppler changes that are not purely motion-related, such as those arising during REM sleep or epileptic seizures. Moreover, rDVARS allows comparison of clutter filtering effectiveness, as it is computed from the power Doppler signal itself. Although we did not use it as such, Power and colleagues suggested that rDVARS can also be computed after denoising to estimate the effectiveness of motion artifact removal (Power et al., 2012).

The complementary properties of these temporal metrics suggest that optimal QC pipelines for fUSI should incorporate multiple measures rather than relying on a single approach. Specifically, GV provides the most direct measure of physical motion, rDVARS effectively captures power Doppler surges independently of slow CBV variations, and rGS offers a familiar metric that aligns with the signal of interest, albeit with limitations in certain experimental contexts. This multi-metric approach allows more nuanced quality assessment that can be tailored to specific experimental designs.

### 4.3 Adaptive clutter filtering improves robustness of awake fUSI

Clutter filtering of raw beamformed ultrasound signals represents the first and arguably most critical step in the fUSI signal processing pipeline, with profound consequences for all subsequent analyses. Our benchmark reveals that optimization at this early processing stage can yield significant improvements in final FC results, potentially eliminating the need for more complex downstream denoising.

#### 4.3.1 Limitations of the SVD clutter filter

Despite its advantages, singular value decomposition (SVD)-based clutter filtering relies on several assumptions that have important implications for fUSI data interpretation. Most fundamentally, these filters assume that clutter and scatterer signals are uncorrelated —an assumption that becomes problematic precisely during awake imaging. When this assumption fails, selecting the correct threshold between clutter and scatterer components becomes challenging.

Moreover, SVD-based clutter filters make interpretation of the power Doppler signal more difficult. Indeed, while frequency-based cutoffs can be directly related to physical properties like scatterer velocity, SVD thresholds lack such straightforward physical interpretations. This ambiguity becomes particularly problematic when estimating CBV, as variations in clutter filtering performance can introduce spurious signal changes that mask true physiological variations. Consequently, while adaptive approaches may more effectively remove clutter during motion, they can paradoxically remove too much signal energy during high-motion periods, producing an apparent drop in global CBV. This contradicts the expected physiological response, where increased movement would typically be associated with increased brain perfusion.

Finally, the mathematical properties of the SVD must be kept int mind. As a least-squares procedure, the SVD is inherently sensitive to outliers, making it vulnerable to spatial variations in clutter intensity. This sensitivity creates reproducibility challenges when applying similar thresholding strategies across different animals or FOV configurations, potentially contributing to the inter-animal variability that has limited fUSI applications.

#### 4.3.2 Applications to fUSI

Our findings suggest that static thresholds for SVD-based clutter filters, although widely used in recent fUSI studies (Brunner et al., 2021; Ferrier et al., 2020; Nunez-Elizalde et al., 2022; Rabut et al., 2020), are suboptimal for awake imaging, as the boundary between clutter and scatterer components fluctuates substantially during acquisition due to animal motion (Baranger et al., 2018). Through direct comparison across four datasets, we demonstrated that adaptive clutter filtering yields significant improvements in motion artifact suppression.

These improvements manifest as increased similarity to group-level FC patterns and reduced inter-animal variability.

The performance of adaptive clutter filters can be understood through their dynamic response to varying motion conditions. Static filters face an inherent compromise: they must use sufficiently high thresholds to adequately suppress clutter during high-motion periods, but this aggressive filtering might inevitably remove relevant signal during low-motion periods. In contrast, adaptive filters can apply more conservative thresholds during calm periods, preserving signals of interest, while dynamically adjusting to more aggressive filtering during motion.

The simple adaptive filter implemented in this study—which maintains approximately constant energy across the FOV or within a brain mask—demonstrates the potential of this approach. However, we acknowledge that this implementation represents just one of many possible adaptive strategies (Baranger et al., 2018). Notably, the constant-energy approach does alter global power Doppler intensity patterns in ways that might complicate interpretation of absolute CBV changes, or the use of the GS as a metric to detect motion. Consequently, this particular implementation should be viewed as a proof of concept for FC analyses, rather than a definitive recommendation.

Our findings highlight the need for continued development of adaptive clutter filtering approaches that balance effective artifact suppression with preservation of physiologically meaningful signals. Future work should evaluate alternative adaptive strategies across diverse experimental paradigms, with particular attention to their effects on different analysis approaches. Advances in this domain have the potential to substantially enhance the reliability and interpretability of fUSI data from awake animals, addressing a critical barrier to wider adoption.

### 4.4 Optimized upstream processing makes scrubbing optional

Previous studies demonstrating the use of fUSI for FC analyses in awake mice have generally applied some form of removal of high-motion volumes (Bertolo et al., 2023; Brunner et al., 2021; Ferrier et al., 2020; Rabut et al., 2020), a process referred to as “scrubbing” in the fMRI literature (Power et al., 2012). Scrubbing is generally performed by removing volumes with significant artifacts from power Doppler time series prior to FC analyses. The main motivation for scrubbing is to remove motion-related variance that is not adequately captured by clutter filters and confound regression methods.

In fMRI, high-motion volumes are generally identified by thresholding metrics such as the framewise displacement (FD) or DVARS (Power et al., 2012; Smyser et al., 2010). Unfortunately, the framewise displacement (FD) metric is not easily applicable to transcranial fUSI data acquired on awake head-fixed mice. Indeed, standard rigid body registration methods for framewise realignment can often fail due to the low SNR and high spatial frequencies of transcranial power Doppler images and the spatial distribution of motion artifacts. These characteristics can make standard gradient descent optimization algorithms unreliable, leading to poor motion correction or even introduction of spurious motion.

Fortunately, we can benefit from well known ultrasound physics principles to estimate the axial velocity of brain tissues, providing an even more direct metric of brain movements (Kasai et al., 1985). In this study, we compared the performance of three scrubbing metrics: the relative global power Doppler signal (rGS), the global axial velocity (GV), and the relative DVARS (rDVARS). Benchmarked strategies included either no scrubbing, or scrubbing by strictly thresholding these metrics.

Our findings indicate that scrubbing effectiveness for FC analyses depends on the metric used to detect motion artifacts. While all scrubbing metrics performed adequately with strict thresholds, rGS-based scrubbing showed limitations. In particular, its use is not recommended with adaptive SVD clutter filters, as these filters may already normalize the GS variations, potentially masking motion events that would be detected using for example rDVARS. In contrast, rDVARS and GV-based scrubbing proved more robust to variations in denoising parameters. The GV metric, in particular, provides a ground-truth measure of brain tissue movement independent of power Doppler signal processing. This makes GV particularly valuable when experimental paradigms are expected to induce genuinely large physiological changes in CBV that might be incorrectly flagged as motion artifacts, such as during task-based experiments. However, other paradigms such as pharmacological manipulations, stroke, or REM sleep may lead to brain-wide CBV changes that are not related to animal motion but may still bias FC analyses. In these cases, rDVARS may be more appropriate, as it allows detection of brain-wide CBV changes that are not related to animal motion.

Interestingly, our results indicate that scrubbing isn’t essential with adaptive clutter filtering and better confound regression methods—for example, aCompCor. Indeed, the best-performing strategies without scrubbing achieved comparable FC similarity to their scrubbing counterparts when using an adaptive clutter filter. This suggests that motion-related variance can be effectively captured at the signal processing level, potentially preserving more of the original acquisition and avoiding the temporal discontinuities introduced by scrubbing. However, it is yet unclear whether the recovered signal still contains relevant neuronal information from motion periods. Thus, further work should be conducted to study the impact of adaptive clutter filters and improved confound regression methods on FC-based metrics. This validation is particularly critical for dynamic FC analyses, where avoiding scrubbing may be beneficial.

In contrast, strategies using static clutter filters showed a clear benefit from scrubbing, and no top-performing strategy used static clutter filters without scrubbing. This result suggests that static clutter filters may not be able to remove motion artifacts sufficiently and require scrubbing of high-motion volumes, and the confound regression methods we tested may not be able to model these artifacts entirely. This finding has important implications for experimental designs where temporal continuity is essential or where motion is correlated with experimental conditions, potentially biasing results if scrubbing is applied.

### 4.5 Band-pass filtering may be detrimental in the presence of motion artifacts

Frequency filtering is a standard component of FC denoising pipelines in both fMRI and fUSI studies (Ferrier et al., 2020; Satterthwaite et al., 2013). However, our results indicate that band-pass filtering in the range (0.01 − 0.1) Hz, which is commonly used in both modalities to isolate frequencies of interest, may be detrimental in the presence of motion artifacts in fUSI acquisitions. Across all datasets, high-pass filtering alone (>0.01 Hz) consistently outperformed band-pass filtering in both motion robustness and FC similarity metrics. Furthermore, all top-performing strategies used high-pass rather than band-pass filtering, regardless of other denoising parameters. This finding has important implications for fUSI denoising pipelines and highlights the need for modality-specific optimizations rather than direct translation of fMRI denoising strategies.

### 4.6 aCompCor confound regression improves control of motion artifacts

Our benchmark revealed that confound regression methods significantly impact the quality of fUSI FC analyses, with aCompCor emerging as the most effective approach across all datasets and denoising parameters. This finding has important implications for fUSI data processing pipelines, which have historically relied heavily on global signal regression (GSR).

Originally developed for fMRI by Behzadi and colleagues (Behzadi et al., 2007), aCompCor derives confound regressors from regions expected to contain minimal neuronal signal of interest (typically white matter and CSF). Unlike GSR, which can remove genuine neural signals that are spatially distributed across the brain (Murphy & Fox, 2017), aCompCor targets physiological noise sources while theoretically preserving signals of interest. In the context of fUSI, where motion artifacts create widespread signal increases that are partially spatially structured, aCompCor’s approach may be particularly advantageous.

Indeed, our results show that aCompCor’s effectiveness for fUSI data persists across different scrubbing approaches and clutter filtering strategies. This consistency suggests that certain noise structures in fUSI data—likely including both physiological noise and motion artifacts —are successfully modeled by aCompCor.

However, it has not escaped our attention that the anatomical definition of noise ROIs for aCompCor remains a critical step. Notably, white matter and CSF masks are particularly thin in our brain slices, and are thus more susceptible to registration errors. Yet, we found that using a degraded white matter + CSF mask significantly lowered the denoising performance (data not shown). Additionally, we benchmarked two additional anatomical masks to derive confound regressors: a random mask and a mask of low-variance voxels, both with as many voxels as the white matter + CSF mask. These masks were included as controls to assess the impact of the anatomical definition of noise regions on denoising performance. aCompCor using the random mask—referred to as random CompCor—performed poorly across all datasets, removing both motion artifacts and signals of interest. On the other hand, aCompCor using the low-variance mask—referred to as low-variance CompCor— performed almost as well as aCompCor when using adaptive clutter filters. This suggests that low-variance voxels may be a suitable alternative to white matter and CSF regions for confound regression. This finding is particularly relevant for studies where white matter and CSF regions are not easily identifiable.

Additionally, we used an arbitrary fixed number of components for aCompCor, which may not be suitable for all datasets. Although we found that changing the number of components from 5 to 20 did not affect significantly the performance of the benchmarked strategies, we noticed that including too many components could lead to removal of relevant signals and degradation of FC patterns. Thus, future work should explore data-driven approaches to determine the optimal number of components for aCompCor.

The marked underperformance of GSR—particularly when used without scrubbing— warrants for caution in its application to fUSI data. Our results suggest that GSR may not be able to effectively model motion artifacts, likely because motion-induced power Doppler surges may have non-uniform spatial distributions. Additionally, our findings indicate that GSR potentially create large spurious anti-correlations that compromise FC analyses. This effect was particularly evident in our evaluation of group-level seed-based maps, where standard strategies employing GSR showed reduced bilateral connectivity and increased spurious correlations compared to aCompCor-based approaches.

Overall, our results collectively suggest that the choice of confound regression strategy for fUSI data should be carefully considered based on the FOV and experimental goals. For most FC analyses in awake mice, our findings strongly favor aCompCor over the traditional GSR approach. Future work should probe further strategies, such as independent component analysis (ICA)-based methods (Griffanti et al., 2014; Pruim et al., 2015), allowing for data-driven extraction of confounds without anatomical priors.

### 4.7 Limitations and perspectives

While our study provides comprehensive guidelines for fUSI denoising in awake animals, several limitations should be acknowledged:

First, the lack of a true ground truth for brain activity limits our ability to definitively assess the biological validity of different denoising strategies. We used group-level FC patterns selected qualitatively by experts as a reference, but these patterns themselves depend on denoising choices. Future studies could incorporate simultaneous electrophysiological recordings or optogenetic manipulations to provide independent validation.

Second, the relationship between motion artifacts and neural activity remains complex. Motion itself can induce neural responses (Steinmetz et al., 2019), making it challenging to fully dissociate artifacts from genuine neural activity. Further research on the neural correlates of movement in head-fixed preparations would help refine artifact identification.

Third, our analysis included relatively few 3D acquisitions (dataset 4), limiting conclusions about volumetric fUSI. As 3D fUSI technology advances, dedicated benchmarks for volumetric data will be necessary to address potential differences in artifact characteristics and optimal denoising strategies.

Fourth, the lack of standardized software frameworks for fUSI denoising remains a significant barrier to reproducibility. Unlike the fMRI community with established tools like AFNI, FSL, SPM, and Nilearn, the fUSI field lacks consensus preprocessing and denoising pipelines. In the present study, we adapted and modified existing fMRI tools to process fUSI data. While offering a practical solution, this approach may not be optimal for fUSI-specific challenges. Further development of dedicated fUSI software tools is needed to facilitate adoption of best practices.

Fifth, while spatial registration across animals is a critical step for extraction of region masks and comparison of group-level FC patterns, registration of transcranial fUSI images remains challenging. Indeed, registration methods for transcranial fUSI remain sparse, with existing studies primarily focused on neuronavigation for probe positioning (Nouhoum et al., 2021). Notably, we used affine registration only, while non-linear registration methods are the norm in neuroimaging (Jenkinson, 2007; Tustison et al., 2021). Unfortunately, transcranial fUSI images display high spatial frequencies, making non-linear registration challenging. Moreover, it is still unclear whether registration of vascular features can lead to proper anatomical alignment. Previous studies have shown that the position of large vessels can vary considerably between mice, making inter-mouse registration of power Doppler images challenging (Bennett & Kim, 2022; Wu et al., 2022). In this study, we mitigated registration errors by combining an iterative registration framework with thorough visual quality control and spatial smoothing. Future work should focus on developing more accurate registration methods specifically tailored to fUSI data, potentially leveraging both power Doppler and B-mode images.

Finally, while our study focused exclusively on transcranial fUSI in head-fixed mice, the physical principles underlying motion artifacts in ultrasound imaging are likely to remain consistent across different experimental preparations and animal models. We anticipate that the methodological principles established here—particularly adaptive clutter filtering, high-pass filtering, and aCompCor confound regression—should generalize to other contexts such as craniotomy preparations, freely-moving setups, larger animal models (rats, ferrets, or non-human primates), and potentially even human applications. The physical interactions between tissue movement, clutter signals, and power Doppler intensity that we characterized are governed by fundamental acoustic principles rather than species-specific factors. However, the magnitude and spatial distribution of these artifacts may vary with anatomical differences, imaging depth, and experimental constraints. Craniotomy preparations may show reduced skull-related artifacts but could introduce new challenges related to brain deformations. Similarly, freely-moving setups could introduce more different motion patterns that might require further optimization of our denoising paradigms. Future studies should systematically validate these denoising approaches across diverse experimental conditions to establish truly generalized protocols for fUSI preprocessing, potentially leading to standardized frameworks that could facilitate cross-species comparisons and translational research.

Despite these limitations, our findings provide valuable practical guidance for researchers using fUSI in awake animals. The benchmarked strategies significantly improve reproducibility and reliability of FC analyses, advancing fUSI as a viable tool for studying brain connectivity in preclinical models.

## 5 Conclusions

This study presents the first comprehensive characterization and systematic benchmarking of motion artifact detection and removal strategies for transcranial fUSI in awake mice. Through detailed analysis of the spatial and temporal characteristics of motion artifacts, we have demonstrated that these artifacts result from complex interactions between tissue motion, signal processing, and potentially physiological responses. Our benchmark of 792 denoising strategies revealed several key methodological principles that significantly improve the quality and reliability of fUSI FC analyses.

First, we established that adaptive clutter filtering substantially outperforms traditional static approaches, especially for FC analyses in awake setups. By dynamically adjusting to varying motion conditions throughout the acquisition, adaptive filters effectively remove motion-related clutter while preserving relevant signals. This finding challenges the current standard practice in the field and offers a path toward more reliable fUSI analyses.

Second, we demonstrated that high-pass filtering alone may be preferable to the commonly used band-pass filtering for fUSI data. This finding contradicts conventional practices borrowed from fMRI and calls for further investigation on the spectral coherence between power Doppler signals and direct neuronal activity.

Third, we identified aCompCor as a superior confound regression method for fUSI data, providing better motion artifact mitigation while preserving relevant FC patterns compared to the widely used global signal regression (GSR). This finding has immediate practical implications for existing fUSI analysis workflows.

Fourth, we showed that with appropriate upstream denoising—particularly adaptive clutter filtering and aCompCor confound regression—scrubbing may become optional for the majority of analyses. This is especially valuable for experimental designs where motion correlates with conditions of interest, making unbiased scrubbing challenging.

Based on these findings, we propose two optimized denoising paradigms tailored to different experimental constraints, with paradigm 1 (using adaptive clutter filtering) demonstrating robust performance across four datasets. These paradigms provide practical recommendations that can be immediately implemented by researchers using fUSI in awake animals.

The methodological advances presented here address a critical barrier to the wider adoption of fUSI for neuroscience research. By substantially improving the reliability and reproducibility of fUSI FC analyses, these techniques enhance the utility of fUSI as a cost-effective, high-resolution tool for studying brain function in preclinical models. Future work should focus on further optimizing adaptive clutter filtering approaches, developing specialized software tools for fUSI data processing, and improving registration methods for inter-subject analyses.

Ultimately, these advances will facilitate more sophisticated applications of fUSI in neuroscience research, enabling studies of complex brain functions and dysfunctions in both health and disease models with unprecedented spatial and temporal resolution.

## 6 Data and Code Availability

The analyses presented in this study were conducted in large parts using a proprietary software currently developed by Iconeus (Paris, France). As such the source code is not publicly available.

The raw datasets supporting the conclusions of this article include raw beamformed I/Q data (>15 TB) and resampled power Doppler data (>1 TB). Given the substantial size of these datasets, they cannot be hosted on standard data repositories. A subset of power Doppler scans denoised using the proposed paradigms is provided on Zenodo for illustration purpose.

Researchers interested in accessing the raw data are invited to contact the Office of Research Contracts at Inserm to initiate discussions regarding data transfer or use.

## 7 Author Contributions

## 8 Funding

Inserm supported this work through the Biomedical Ultrasound Technology Research Accelerator (ART).

## 9 Declaration of Competing Interests

Z.L, B-F.O., and T.D. are co-founders and shareholders of Iconeus and have received funding from Iconeus for research on functional ultrasound imaging. Some of their patents are licensed to Iconeus company. B-F.O., Z.L., and T.D. are members of the scientific board of Iconeus company. S.L.M-D., F.C.P., and B-F.O. are employees of Iconeus company. Z.L. and T.D. are consultants for Iconeus Company. The remaining authors declare no competing interests.

## 10 Acknowledgements

The authors thank Luc Eglin for valuable discussions about the methods used in this study.

## 11 Supplementary Material

**Supplementary Figure 1:**
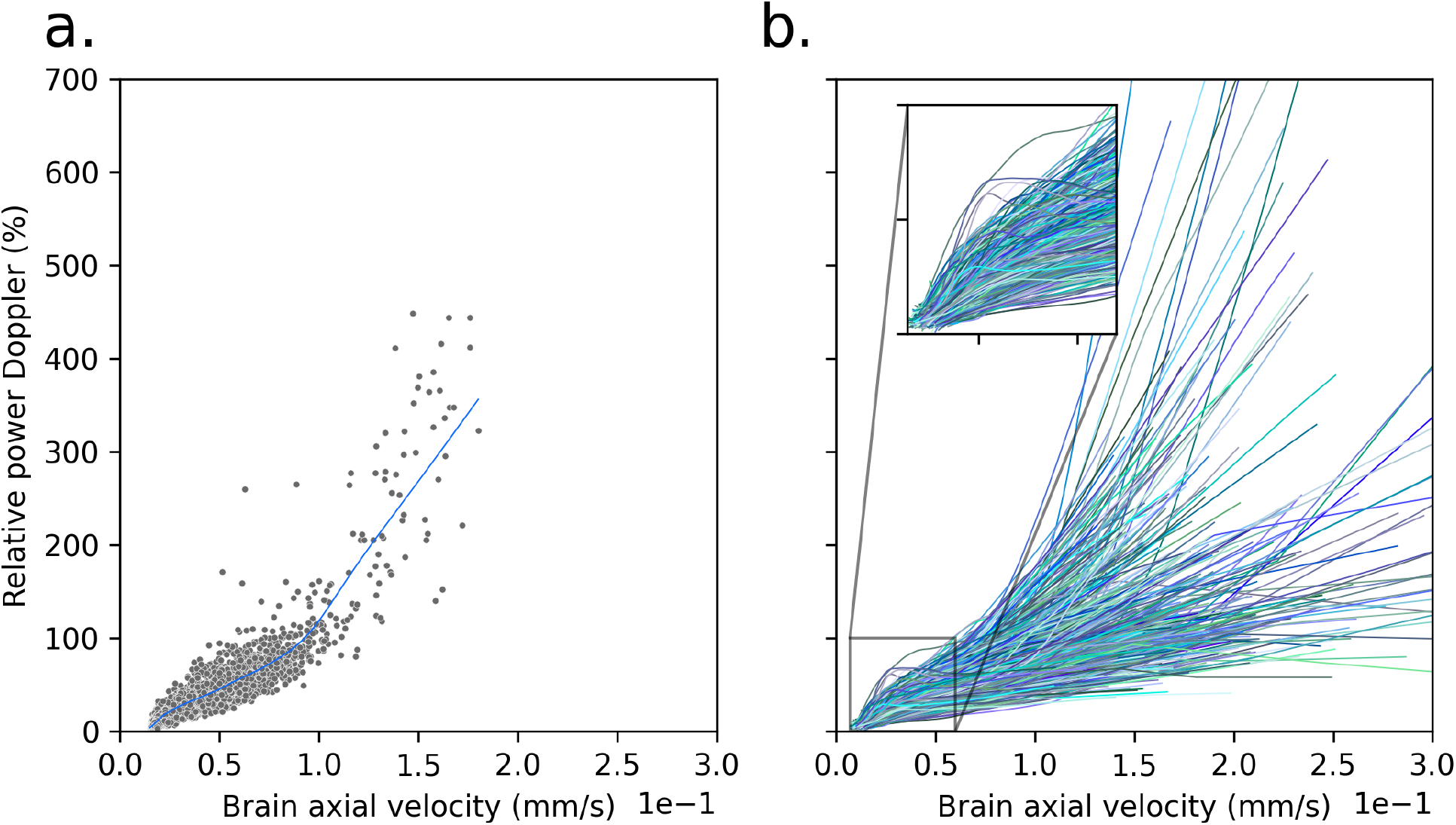
Power Doppler intensity is related to global axial velocity (GV). (**a**.) The relative power Doppler signals averaged over brain voxels is plotted against the GV. A locally weighted scatterplot smoothing (LOWESS) curve is fitted on the data. (**b**.) LOWESS curves for all acquisitions from datasets 1 and 2 are shown, showing a non-linear dependence between power Doppler intensities and GV, even for small velocities.

**Supplementary Figure 2:**
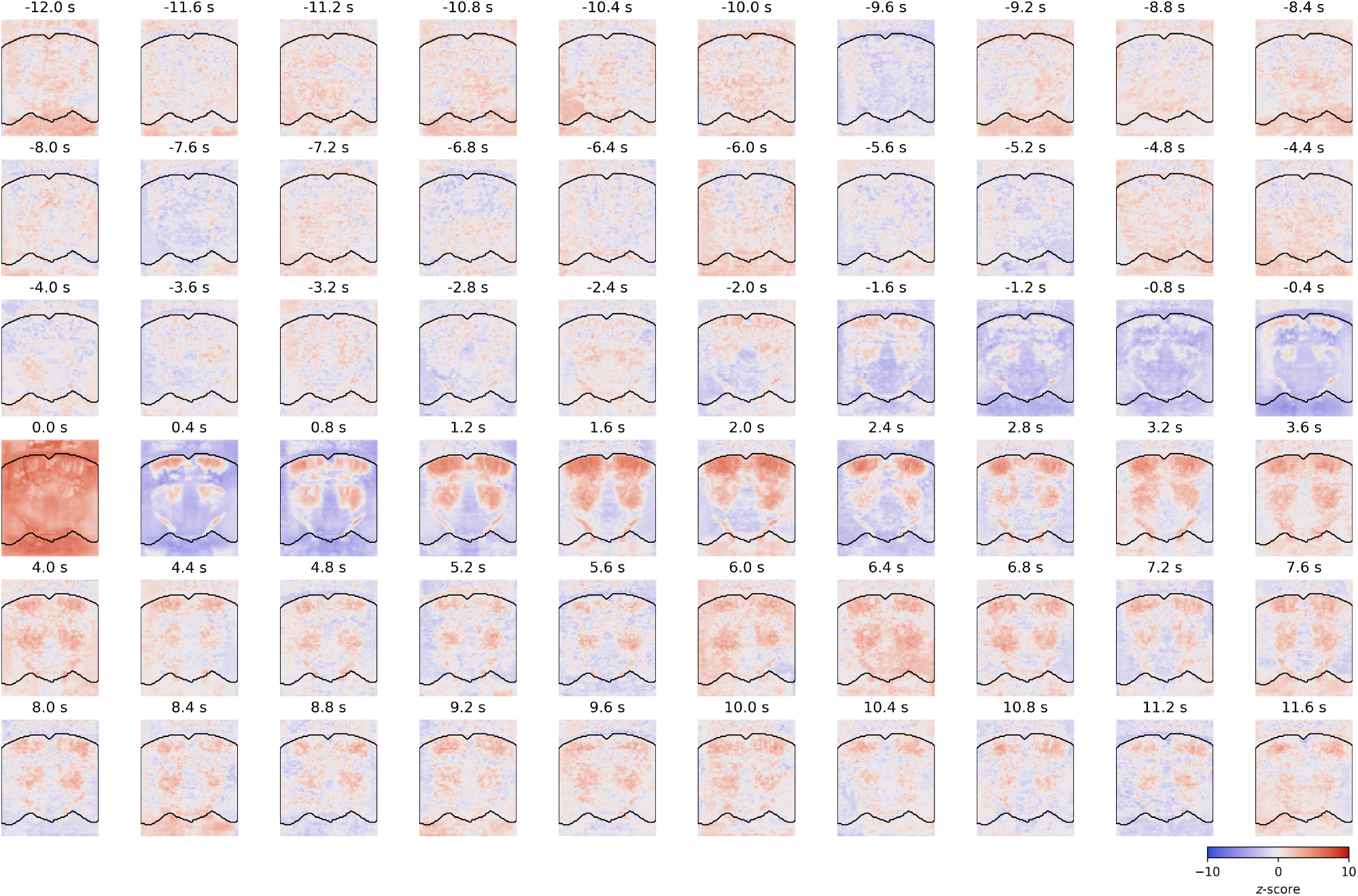
Brain movement induces both immediate and delayed effects on power Doppler signals. Maps are shown for all delays for the FIR model in Figure 2.

**Supplementary Figure 3:**
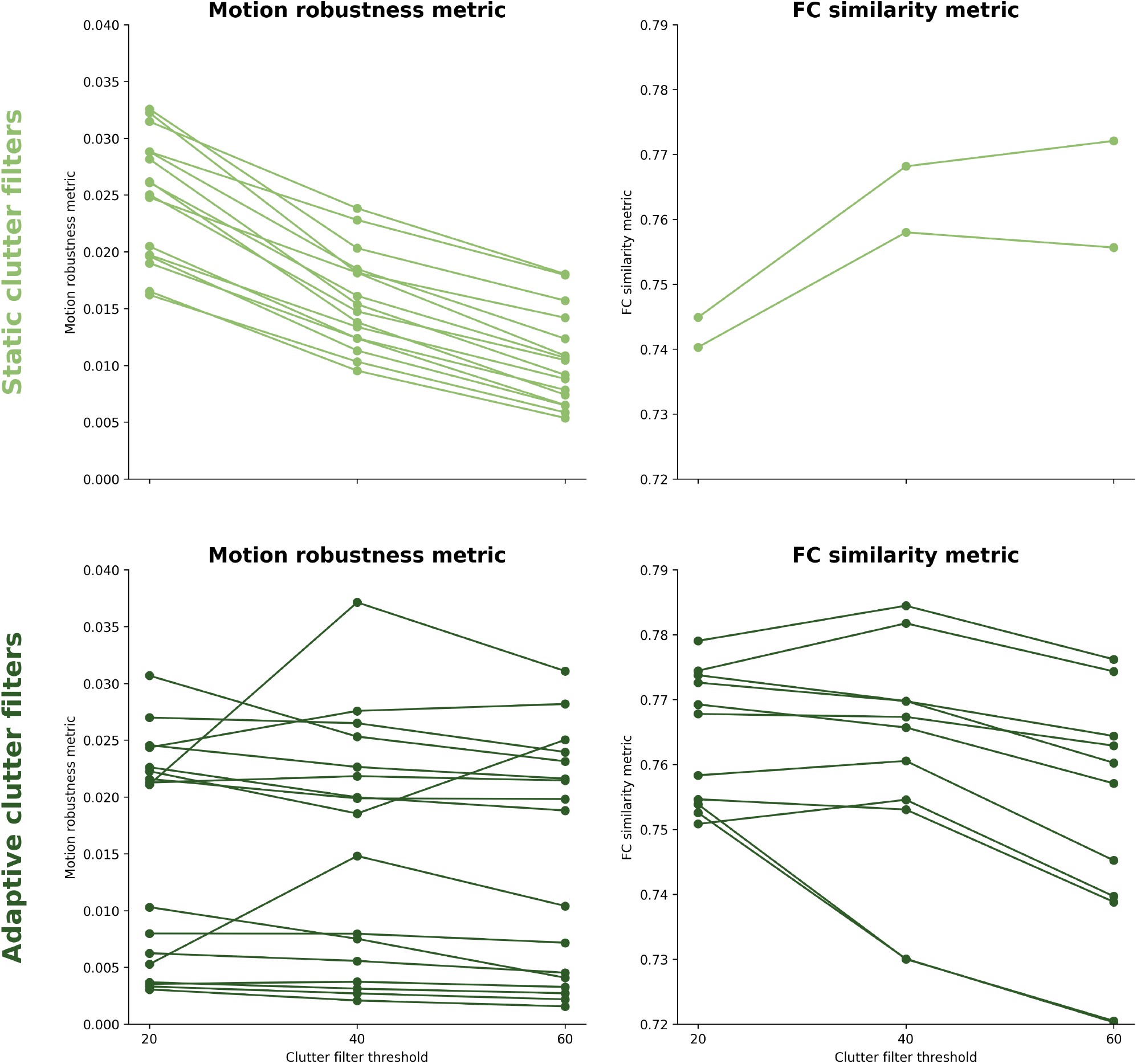
Metrics across clutter filter thresholds. Slope graphs show the motion robustness (**left**) and FC similarity (**right**) metrics across clutter filter thresholds. Metrics are shown using dataset 1 for all strategies that pass the corresponding metric threshold (either MSE < 0.025 or Dice > 0.75). Each row shows The top row shows the metrics for strategies using static SVD*T* clutter filters, while the bottom row shows the metrics for strategies using adaptive SVD*T* clutter filters.

**Supplementary Figure 4:**
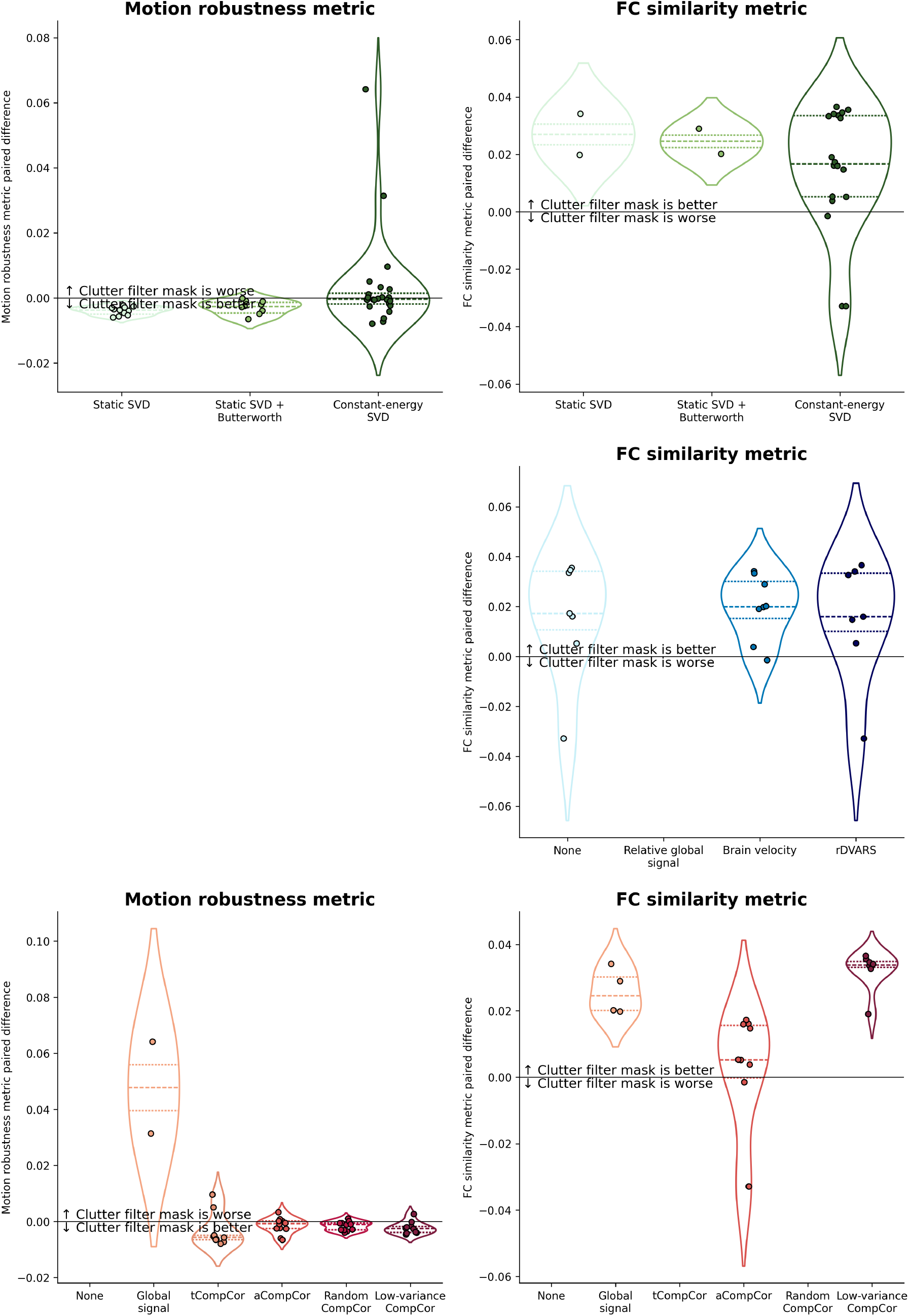
Paired metric differences for clutter filter masking strategies. Violin plots show the paired differences in motion robustness (**left**) and FC similarity (**right**) metrics between clutter filter masking strategies. Paired differences are computed using dataset 1 for all strategies that pass the corresponding metric threshold (either MSE < 0.025 or Dice > 0.75). Each row shows the same paired differences, colored by clutter filter, scrubbing metric, and confound regression method.

**Supplementary Figure 5:**
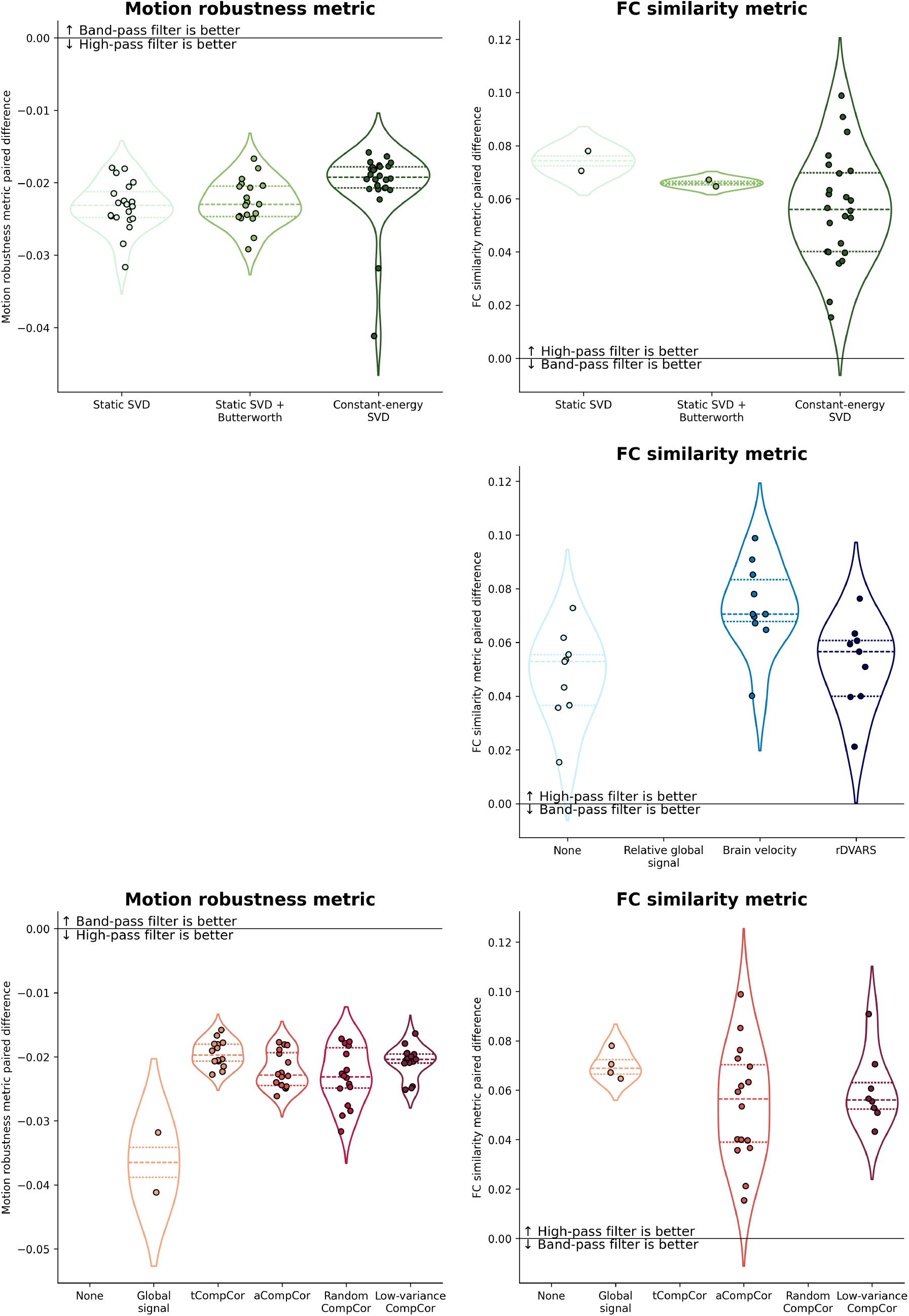
Paired metric differences for frequency filtering strategies. Violin plots show the paired differences in motion robustness (**left**) and FC similarity (**right**) metrics between high-pass and band-pass filtering strategies. Paired differences are computed using dataset 1 for all strategies that pass the corresponding metric threshold (either MSE < 0.025 or Dice > 0.75). Each row shows the same paired differences, colored by clutter filter, scrubbing metric, and confound regression method.

